# Multimodal profiling of peripheral blood identifies proliferating circulating effector CD4^+^ T cells as predictors for response to integrin α4β7-blocking therapy in patients with inflammatory bowel disease

**DOI:** 10.1101/2023.10.01.560386

**Authors:** Veronika Horn, Camila Cancino, Lisa Steinheuer, Benedikt Obermayer, Konstantin Fritz, Anke L. Nguyen, Christina Plattner, Diana Bösel, Marie Burns, Axel Ronald Schulz, Eleni Mantzivi, Donata Lissner, Thomas Conrad, Mir-Farzin Mashreghi, Elena Sonnenberg, Dieter Beule, Lukas Flatz, TRR241 IBDome Consortium, Zlatko Trjanoski, Carl Weidinger, Henrik E. Mei, Britta Siegmund, Kevin Thurley, Ahmed N. Hegazy

## Abstract

Despite the success of biological therapies in inflammatory bowel disease (IBD), patient management remains challenging due to a lack of therapy response predictors. Here we prospectively sampled two cohorts of IBD patient cohorts receiving the anti-integrin α4β7 antibody vedolizumab. Samples were subjected to mass cytometry, single-cell RNA sequencing, single-cell V(D)J sequencing, serum proteomics, and multidimensional flow cytometry to comprehensively assess vedolizumab-induced immunological changes in the peripheral blood and their potential associations with treatment response. Vedolizumab induced changes in the abundance of both circulating innate and adaptive immune cell compartments and modified the T cell receptor diversity of circulating gut-homing CD4^+^ memory T cells. Through integration of multimodal parameters and machine learning, we identify that pretreatment activated proliferating CD4^+^ memory T cell abundance is associated with treatment failure, independent of clinical variables, thereby providing a reliable predictive classifier with significant implications for the personalized management of IBD patients.

## MAIN

Inflammatory bowel disease (IBD) is a chronic, relapsing inflammatory gastrointestinal disease driven by the complex interplay between host genetics, environmental triggers, microbial dysbiosis, and intestinal immune dysregulation^1–4^. Biologic therapies have improved outcomes for IBD patients, and one such biologic, vedolizumab, disrupts interactions between MAdCAM-1, expressed by intestinal endothelial cells, and integrin α4β7, present on circulating leukocytes to sequester activated gut-homing cells within the circulation^5–9^. Vedolizumab effectively induces and maintains remission in patients with Crohn’s disease^10^ and ulcerative colitis^11^, particularly those with colonic inflammation, with a remarkable safety profile. However, only a fraction of patients respond to vedolizumab and many do not achieve complete remission, necessitating a “trial-and-error” approach^12,13^. There is a clear unmet need to decipher the immunological changes in specific patient populations to classify treatment response^14,15^. We therefore employed a multimodal profiling approach to investigate vedolizumab-induced immunological changes in a prospective cohort of IBD patients receiving vedolizumab and age- and sex-matched non-IBD controls (**Fig. 1a, b**, and **Extended Data Fig. 1a, Suppl. Table S1**). Clinical response was assessed, and stool calprotectin levels faithfully discriminated responders from non-responders (**Extended Data Fig. 1b-e**). To comprehensively capture changes in circulating immune cell composition and the inflammatory milieu, we profiled peripheral blood using high-throughput mass cytometry (CyTOF), flow cytometry (FACS), and serum proteomics (Olink) (**Fig. 1b**). In a sub-cohort, we performed single cell proteogenomics (CITE-Seq), single cell (sc)RNA-seq, and immune repertoire profiling on sorted CD45^+^ cells to investigate integrin α4β7 expression and changes in T and B cell receptor (TCR/BCR) repertoires upon vedolizumab treatment.

**Figure 1.**
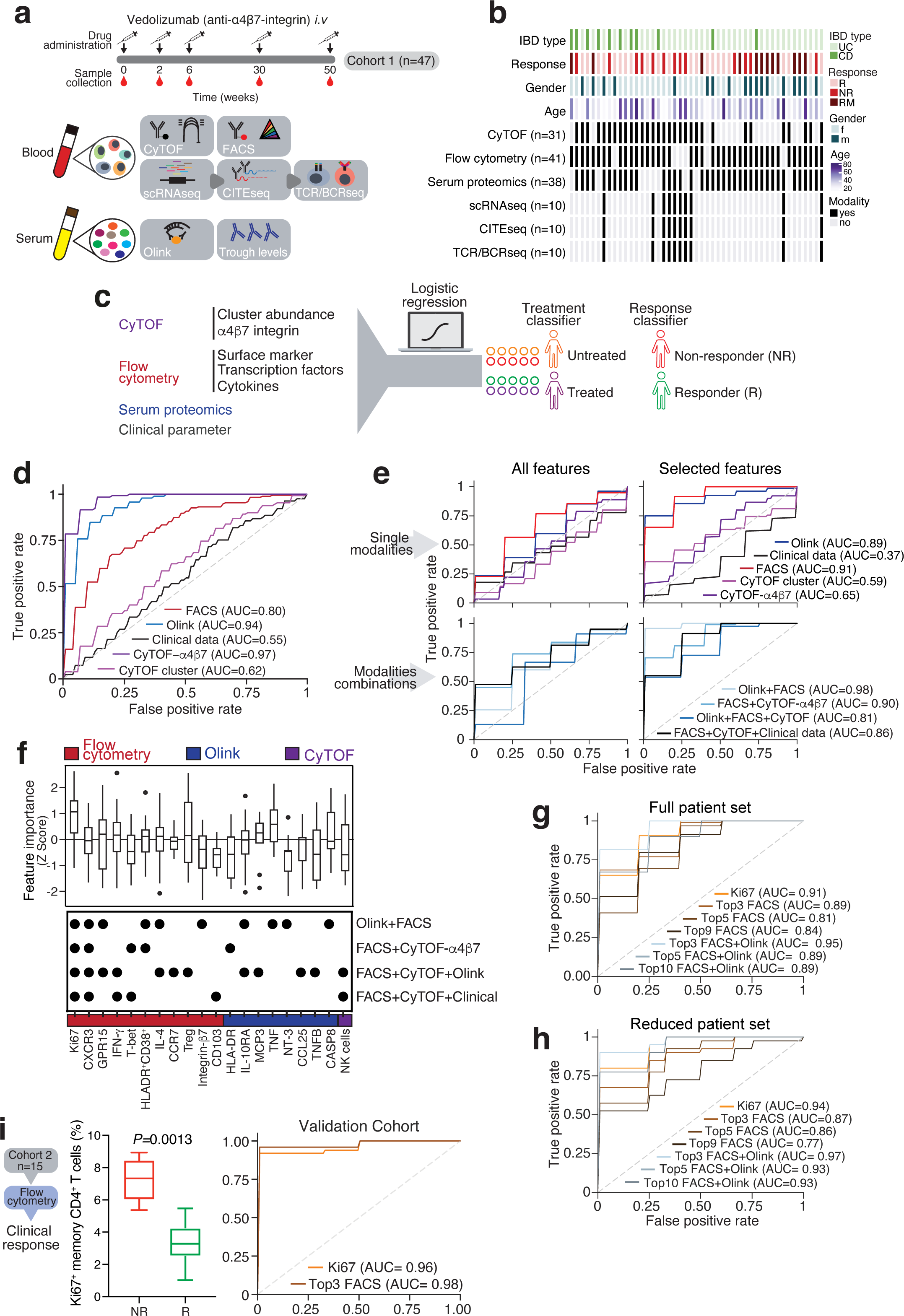
Machine learning-based classification identifies vedolizumab-induced immune changes and classifies therapy response. (a) Study design. (b) The study cohort (n=47) and analyses applied to each IBD patient sample. (c) Machine learning approach used to identify the effects of vedolizumab treatment on circulating immune cells and to classify treatment response. (d) Capacity of clinical parameters, serum proteomics, mass cytometry, and flow cytometry data to display vedolizumab treatment signatures shown as receiver operating characteristic curves (ROC) with corresponding area under the curve (AUC) values. (e) Predictive ability of individual data sets (top) and data combinations showing the highest AUC values (bottom) to classify vedolizumab response. Classifications using all features (left) and the top 10% most predictive features for Olink, FACS, CyTOF, CyTOF-α4β7 from the ranked logistic regression coefficients (right). (f) Overview of the 19 most predictive features for vedolizumab response. Feature importance based on Z-score-transformed model coefficients from the most predictive classification models (top) and the frequency of each predictor across the best performing models (bottom). (g) Predictive ability of combinations of FACS and Olink+FACS markers with the highest feature importance. Models were trained using all patients included in the respective data modalities. (h) Predictive capacity of selected FACS and Olink+FACS marker panels shown in (f). Models were trained on 15 randomly selected patients included in the Olink-FACS panel. (i) Biomarker validation in an independent cohort of IBD patients treated with vedolizumab (n=15). Shown is the percentage of Ki67^+^ within memory CD4^+^ cells (left bar graph) and the predictive capacity using Ki67 or the top three FACS markers (Ki67, CXCR3, GPR15), as indicated. Mann-Whitney test.

We identified 18 distinct circulating cell populations by CITE-Seq and 16 by CyTOF (**Extended Data Fig. 2**). scRNA-seq analysis demonstrated widespread *ITGA4* and *ITGB7* expression in major circulating immune cell lineages (**Extended Data Fig. 3a-d**). CyTOF confirmed broad integrin α4β7 protein expression in T cells, monocytes, eosinophils, basophils, B cells, and NK cells, with slight differences in integrin α4β7 expression observed between healthy controls and IBD patients (**Extended Data Fig. 3e-h**). Using single-cell V(D)J sequencing to examine changes in circulating T and B cell compartments (**Extended Data Fig. 4a, b and 5a, f**), we observed a significant increase in clonal diversity in CD4^+^ memory T cells but not in CD8^+^ T and B cell subsets following vedolizumab treatment (**Extended Data Fig. 4c, 5b-d, g**). Specifically, there was an enhanced TCR diversity in central and effector memory CD4^+^ T cells as well as CD4^+^ T cells expressing *ITGA4* and *ITGB7* (**Extended Data Fig. 4d-f**), but not in other gut-homing CD4^+^ T cell subsets (GPR15^+^ or integrin αE^+^; **Extended Data Fig. 4e**). Consistent with these findings, the abundance of circulating memory CD4^+^ T cells (but not CD8^+^ T and B cells) increased after vedolizumab treatment (**Extended Data Fig. 4g-i, and Extended Data Fig. 5e, h, i**). We focused our flow cytometry analysis on CD4^+^ memory T cells given the observed alterations in their diversity and abundance.

Next, we applied machine learning to our multimodal dataset to examine the influence of vedolizumab on circulating immune cells and to investigate potential predictive classifiers of vedolizumab response (**Fig. 1c, Suppl. Table 2**). Using a logistic regression classifier, vedolizumab most altered integrin α4β7 abundance within circulating immune cells (CyTOF-α4β7; AUC=0.97; **Fig. 1d and Extended Data Fig. 6a-c**) and serum protein levels (Olink; AUC=0.94; **Fig. 1d and Extended Data Fig. 6d, e**). The ten most altered features within each dataset were the abundance of integrin α4β7^+^ eosinophils and cDC populations, memory CD4^+^ T cells (**Extended Data Fig. 6b, c**), and serum markers including eotaxin (CCL11), delta Notch-like epidermal growth factor-related receptor (DNER), tumor necrosis factor B (TNFB), and CCL28 (**Extended Data Fig. 6d, e**).

We applied different classification methods on all features within each dataset to identify treatment response classifiers (**Fig. 1c, Extended Data** Fig. 7a), but this approach did not classify treatment response (**Fig. 1e**, upper left panel, **Suppl. Table 3**). Due to comparable classification performance (**Extended Data Fig. 7a**), logistic regression was used to explore combinations of different data modalities to improve classification performance, but again this did not classify response (**Fig. 1e**, lower left panel, **Suppl. Table 3**). We then explored a reduced feature set by selecting the top 10% most predictive features from the ranked logistic regression coefficients, which significantly improved classification performance using Olink and FACS data (**Fig. 1e**, upper right panel; **Extended Data Fig. 7b**).

Utilizing the top 10% most predictive features from Olink and FACS data together enhanced the classification performance, yielding an AUC of 0.98 (**Fig. 1e**, lower right; **Extended Data Fig. 7c, Suppl. Table 3**). The classification performance remained convincing when using data from only 15 patients profiled with all modalities (**Extended** Data Fig. 7d). The most predictive features for classifying treatment response included parameters related to CD4^+^ memory T cells, such as Ki67, chemokine receptors, cytokines, serum inflammatory markers, and NK cell abundance (**Fig. 1f**, and **Extended Data Fig. 7e, f**). Indeed, using the three key features (Ki67, GPR15, TNF) from FACS and Olink data, we were able to predict vedolizumab response with an AUC of 0.95 (**Fig. 1g**). Notably, the predictive ability of Ki67^+^CD4^+^ T cells alone yielded an AUC of 0.91 (**Fig. 1g**). Assessing the robustness of the identified markers, using only 13 patients, the classification performance remained compelling (**Fig. 1h**).

To validate the identified features for therapy classification, we evaluated the top three predictive FACS features (Ki67, GPR15, CXCR3) in an independent validation cohort, which achieved an AUC of 0.98 (**Fig. 1i**). Moreover, the abundance of Ki67^+^CD4^+^ memory T cells accurately classified treatment response with an AUC of 0.96 (**Fig. 1i**). Ki67 expression was only increased in CD4^+^ memory T cells but not in CD8^+^ or regulatory T cells (**Extended Data Fig. 8a, b**). There were significantly more Ki67^+^ CD4^+^ memory T cells in IBD patients before treatment than in healthy controls, and these remained high even at six weeks (**Extended Data Fig. 8c**).

To gain further insight into the characteristics of Ki67^+^ memory CD4^+^ T cells, we performed CITE-Seq on newly sorted CD3^+^CD4^+^CD45RA^-^ memory T cells from the same donors used in our initial PBMC scRNA-seq profiling (**Fig. 1b**). Twelve distinct CD4^+^ memory T cell subsets were identified by hierarchical clustering (**Fig. 2a, Extended Data Fig. 9a-c**). Notably, Ki67^+^ memory CD4^+^ T cells were predominantly enriched in cluster 1 (**Fig. 2b, c**), which was significantly increased in non-responders before treatment (**Fig. 2d**).

**Figure 2.**
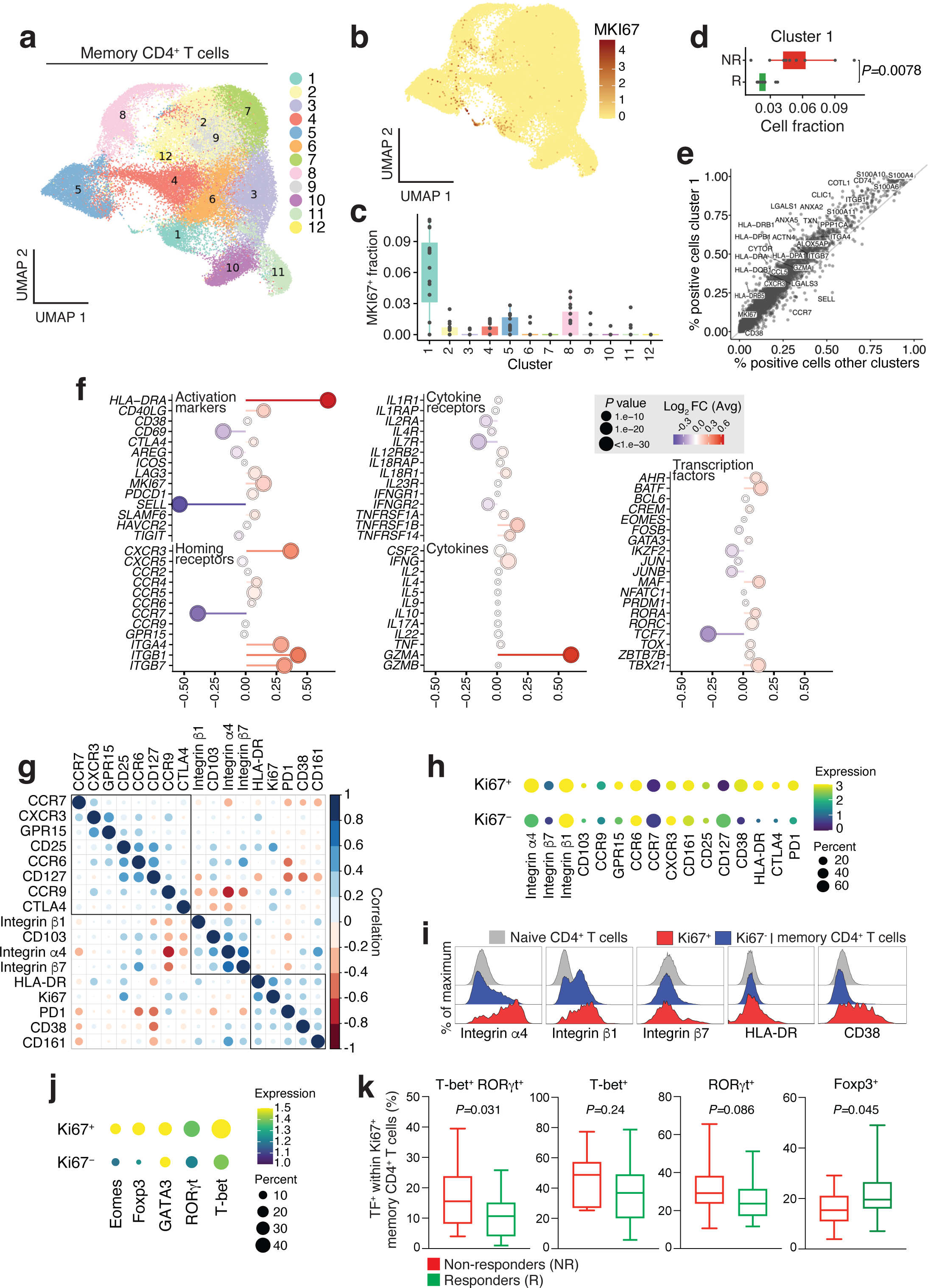
Characteristics of proliferating memory CD4^+^ T cells before treatment. (a) UMAP plot of CD4^+^ memory cells using combined PBMC and CD4^+^-sorted scRNA-seq data from 5 HC and 10 IBD patients. (b) Normalized expression of *MKI67* in CD4^+^ memory cells. (c) Fraction of *MKI67*^+^ cells within different clusters. (d) Fraction of *MKI67*^+^ cells in cluster 1 for responders (R) and non-responders (NR), respectively. Mann Whitney test. (e) Scatter plot of marker genes for cluster 1 showing percentage of cells with detectable expression in cluster 1 compared to all other clusters (f) Lollipop graph illustrating fold expression of selected genes in cluster 1 compared to other clusters. The color scale represents the average log2-fold expression, and the dot size represents the *P*-value. Genes are categorized according to different functions, including activation markers, cytokine receptors, transcription factors, cytokines, and homing receptors. (g) Dot plot showing correlations between FACS surface markers within CD4^+^ memory T cells in IBD patients before therapy (n=41). (h) Dot plot showing geomean expression of FACS surface markers on Ki67^+^ or Ki67^-^ memory CD4^+^ T cells in IBD patients before therapy (n=41). Expression normalized to naïve CD4^+^ T cells. (i) Histograms of FACS surface marker expression in a representative IBD patient. Shown is the expression of the indicated markers on naïve, Ki67^+^, and Ki67^-^ memory CD4^+^ T cells. (j) Dot plot showing geomean expression of FACS transcription factors on Ki67^+^ or Ki67^-^ memory CD4^+^ T cells in IBD patients before therapy (n=41). Expression is normalized to naïve CD4^+^ T cells. (k) Expression of the indicated transcription factors within Ki67^+^ and Ki67^-^ memory CD4^+^ T cells in vedolizumab responders (R) vs. non-responders (NR). n=18-32. Mann Whitney test.

Next, we examined the characteristics of cluster1 and assessed the expression of a set of classical activation and homing markers, cytokine receptors, cytokines, and T cell-related transcription factors (**Fig. 2e, f, Suppl. Table 4**). Cluster 1 revealed signs of activation and proliferation, with upregulation of *HLADRA, CD40LG,* and *MKI67* and downregulation of *IL7R, and SELL* (**Fig. 2f, Extended Data** Fig. 8d; activation markers and cytokine receptors). Furthermore, cells in cluster 1 showed high *ITGA4, ITGB1*, and *ITGB7* levels and expressed *CXCR3* and *CCR6* and expressed *IL12BR2, and IL18R1,* and showed down-regulation of *IFNGR2* (**Fig. 2f**; homing and cytokine receptors). Several transcription factors were upregulated, including TBX21, RORC, MAF, and TOX (**Fig. 2f**; transcription factors).

Next, we used multidimensional flow cytometry to confirm the CITE-Seq results and to explore the functionality of the predictive Ki67^+^ effector CD4^+^ memory T cell signature. There was a positive correlation between Ki67 and HLA-DR, CD38, and PD-1 in memory T cells (**Fig. 2g**), particularly CD38-HLA-DR double-positive cells (**Extended Data Fig. 8e**). Ki67^+^ CD4^+^ memory T cells showed increased expression of CD38, HLA-DR, CTLA4, and PD-1 and decreased expression of CD127 (**Fig. 2h, i, Extended Data Fig. 8f, g**). Nearly 80% of the Ki67^+^ CD4^+^ memory T cells expressed integrin α4β1, and only 20% of these cells expressed integrin α4β7 (**Fig. 2h, i, Extended Data** Fig. 8f, g). Ki67^+^ CD4^+^ memory T cells were enriched in CXCR3, and GPR15, both of which were associated with treatment outcome (**Fig. 1f and 2h, i, Extended Data Fig. 8f, g**). Indeed, inflamed intestinal tissue in IBD patients showed elevated expression of the corresponding chemokine ligands and adhesion molecules (**Extended Data Fig. 10**).

Finally, we further explored the characteristics of Ki67^+^ CD4^+^ memory T cells, which showed increased expression of the Th1- and Th17-lineage transcription factors T-bet and RORγt, as well as Eomes (**Fig. 2j, Extended Data Fig. 8h)**. This was consistent with the observed expression of IFN-γ and IL-17A in Ki67^+^ CD4^+^ memory T cells (**Extended Data Fig. 8i**). Lastly, we observed an increased co-expression of T-bet and RORγt and a reduction in FoxP3 in non-responders compared with responders (**Fig. 2k**).

Our analysis of the impact of anti-integrin α4β7 treatment on the circulating immune cell landscape in IBD revealed four main observations. First, there was a broad expression of integrin α4β7 in circulating immune cell types, which could explain the changes in the abundance of various immune cells after vedolizumab. Second, vedolizumab increased TCR repertoire diversity in α4β7^+^ CD4^+^ memory T cells. Third, by integrating these features, we could classify response to vedolizumab before therapy initiation. Finally, we confirmed that circulating proliferating effector memory CD4^+^ T cells are a candidate predictor of vedolizumab failure in IBD patients. Thus, despite the limitations of a relatively small sample size, most patients having ulcerative colitis, and potential bias because not all patients were profiled using all platforms, our study comprehensively characterized the circulating immune cell landscape and identified predictive factors for treatment response.

The combination of multimodal profiling and machine learning techniques allows the generation and analysis of large datasets to identify the immunologic impact of therapeutic interventions and enables the unbiased detection of patterns that may classify therapeutic outcomes. Currently, there are currently no blood-based predictive biomarkers for IBD patients^14–16^. Our proposed biomarkers (Ki67, GPR15, and CXCR3) can be easily measured in peripheral blood before treatment using standard flow cytometry, making them clinically accessible. Ki67 expression in CD4^+^ memory T cells alone showed excellent predictive power and could be easily measured in routine diagnostic settings. The precise role of cycling CD4^+^ memory T cells in intestinal inflammation in vedolizumab non-responders remains unexplored. These cells predominantly express integrin α4β1, making them unresponsive to vedolizumab. Exploring the impact of integrin α4 antagonists that target these cells may be of benefit to IBD patients who are resistant to vedolizumab^6,17^.

## MATERIAL AND METHODS

### Study design and approval

A total of 47 patients (cohort 1; **Suppl. Table 1**) and 15 patients (cohort 2; **Suppl. Table 5**) receiving Vedolizumab at the outpatient and inpatient clinic of the Department of Gastroenterology, Infectiology, and Rheumatology at Charité University Medicine Berlin were enrolled in either a prospective study (VEPREDEX#EA4/162/17) or a prospective biobank (IBDome-study; EA4/162/17) which were approved by the Charité University Medicine Berlin ethics committee. Donors in the first cohort were recruited between July 2018 and November 2021, while patients for the second cohort were included from December 2021 to February 2023. We also included 41 age- and sex-matched healthy donors (HC) for comparison. All donors provided informed written consent for their participation in the study. In addition to bio-sampling, we collected relevant (para-) clinical data, including Harvey-Bradshaw-Index (HBI)/Partial Mayo Score (PMS), C-reactive protein levels, and leukocyte and thrombocyte counts. Responders were classified as patients who achieved a minimum reduction of three points in the HBI and at least two points in the PMS after 30 weeks of vedolizumab treatment.

### Sample collection and banking

Peripheral blood samples were collected from both IBD patients and healthy donors (HD) at defined time points before and during therapy (**Figure 1A**) using heparin and serum tubes. Samples were collected before the first infusion of vedolizumab (week 0), and before the following infusions (week 2, or 6: before the 2^nd^ or 3^rd^ infusion at week 2 or 6). For peripheral blood mononuclear cells (PBMCs) isolation, peripheral blood was diluted 1:1 in 1x PBS and subjected to density gradient centrifugation using Pancoll human (PAN-Biotech, Aidenbach, Germany; P06-1391500) at 800 x g for 20 minutes at room temperature. The resulting PBMC fraction was then resuspended in a freezing medium composed of 90% FCS and 10% DMSO. Subsequently, the PBMCs were frozen using a Cryo-freezing container (Nalgene, Rochester, NY) and stored in liquid nitrogen for long-term preservation. A proteomic stabilizer PROT-1 (Smart Tube Inc., Las Vegas, NV) was used for whole blood cryopreservation according to the manufacturer’s instructions, and cryopreserved whole blood samples were stored at −80°C. Patient serum samples were obtained by centrifugation at 2000 x g for 7 minutes at room temperature. The resulting serum was immediately frozen at −80°C for subsequent analysis.

### PBMC thawing for stimulation

Frozen PBMCs were thawed in a 37°C water bath and then transferred to a preheated thawing medium consisting of RPMI 1640 Medium GlutaMAX™ Supplement, 10% FCS, 1% penicillin/streptomycin, 50 µM 2-mercaptoethanol, and 50 U/mL DNase I (Sigma-Aldrich, St Louis, MO). Cells were centrifuged at 350 x g for 10 minutes at room temperature. The cells were then resuspended in RPMI 1640 Medium GlutaMAX™ Supplement, 10% FCS, 1% penicillin/streptomycin, and 50 µM 2-mercaptoethanol, counted, and 1 million cells were used for flow cytometry.

### Stimulation of PBMCs with PMA/Ionomycin

Thawed PBMCs were incubated with 100 µl of a stimulation mixture containing 5 ng/ml PMA (Sigma-Aldrich, St Louis, MO; P1585-1MG) and 500 ng/ml ionomycin (Sigma-Aldrich, St Louis, MO; I0634-1MG,) at 37°C for 30 minutes. After stimulation, 50 µl of brefeldin A at a concentration of 5 µg/ml (Cayman Chemical, Ann Arbor, MI; Cay11861-25) was added, and the cells were incubated for a further 3.5 hours at 37°C. After a total of four hours of stimulation, the cells were harvested for intracellular cytokine (IC) staining.

### Flow cytometry

PBMCs were stained with surface antibodies in FACS buffer (1x PBS, 0.05% BSA, 0.01% NaN3, 2 mM EDTA; **Suppl. Table 6**) for 30 minutes. Cells were then fixed and permeabilized with the eBioscience™ Foxp3/Transcription Factor Staining Buffer Set (Invitrogen, Waltham, MA) to allow intracellular staining for Ki67, and transcription factors. For intracellular cytokine staining, PBMCs were fixed with 1x BD FACS Lysing Solution (BD Biosciences, Franklin Lakes, NJ) following stimulation with PMA/ionomycin, and surface staining, and live/dead staining (**Suppl. Table 6**). Subsequently, intracellular cytokines were stained after permeabilization with 0.05% saponin (Sigma-Aldrich). The gating strategy to identify CD4^+^ memory T cells and the expression of various surface markers and staining controls are shown in Technical supplementary figure 1, 2 and 3. Cell suspensions were supplemented with Precision Count Beads (BioLegend, San Diego, CA) and then acquired on a BD FACSymphony (Configuration 5B 8V 3R 5YG 7UV). Daily quality control was performed using Sphero Rainbow Calibration Particles (BD Biosciences, Franklin Lakes, NJ). Flow cytometry data was analyzed using FlowJo software (BD, V10.4), and subsequent analyses were performed using R and Prism v8 (GraphPad Software, La Jolla, CA). Clinical information regarding the patients included in the flow cytometry analysis can be found in **Suppl. Tables 5 and 7.**

### Cellular barcoding and mass cytometry

Frozen whole blood samples were thawed, stained, and acquired in batches of 15 samples that included all time points from a patient and matched healthy controls. An anchor sample was included in each run to control for batch effects. The thawing process and mass cytometry staining were performed following as previously described^18^. After thawing and erythrocyte lysis, total cell counts were obtained using a MACSquant cytometer. Subsequently, 1.5 million cells per sample were barcoded using the Cell-ID Palladium Barcoding Kit (Standard BioTools, South San Francisco, CA) according to the manufacturer’s instructions and pooled into a single batch. To minimize nonspecific binding, cells were incubated with 0.2 mg/ml Beriglobin (CSL Behring, King of Prussia, PA) for 10 minutes to block Fc receptors. After washing, the cell suspensions were treated with heparin (Ratiopharm, Ulm, Germany) at a concentration of 100 U/ml for 15 minutes to reduce eosinophil background artifacts. Cell surface staining was performed for 30 minutes in the presence of heparin using the antibody panel listed in **Suppl. Table 8**. After washing, cells were incubated with the appropriate secondary antibody for 15 minutes at room temperature. Antibodies were either pre-labeled (Standard BioTools), labeled in-house with platinum, or labeled using MAXPAR X8 labeling kits (Standard BioTools) according to the manufacturer’s instructions. Antibody cocktails were prepared in advance and thawed immediately prior to staining.

After staining, cells were washed and suspended in 4% paraformaldehyde (Electron Microscopy Sciences, Hatfield, PA) overnight. The next day, cells were washed again and counterstained with an iridium-based DNA intercalator (Standard BioTools). Cell suspensions were repeatedly washed with deionized water, passed through a 30 µm cell strainer (Corning, Corning, NY), and spiked with 10% (v/v) EQ Four Element Calibration Beads (Standard BioTools). Daily quality control and adjustment of the Helios mass cytometer (Standard BioTools) were performed prior to acquisition. A total of 300,000 events per sample were acquired at a rate of 200-300 events per second. The samples were acquired in two cohorts and normalized to account for any batch-dependent effects. Clinical information regarding the patients included in the mass cytometry analysis can be found in **Suppl. Table 9**.

### Mass cytometry data processing and algorithm-based high-dimensional analysis

Normalization to the spiked-in beads was performed immediately after acquisition using Fluidigm software (Standard BioTools) based on EQ Four Element Calibration Beads (passport P13H2302). Archsinh transformation with cofactor = 5, debarcoding and matrix-based compensation were performed in the R environment using the CATALYST workflow. Events were then manually pre-gated and gate-cleaned in FlowJo with the exclusion of neutrophils, as shown in (Technical supplementary figures 4). In all conducted experiments, we acquired 13,311,287 cells derived from 154 samples, which were subsequently analyzed. Marker expression between the two cohorts was normalized to the 95th percentile using the batch adjust function in R^19^. The resulting data set showed no significant difference between the two cohorts (**Technical Supplementary** Figure 5).

Subsequent algorithm-based high-dimensional analysis was performed on the High Performance Computing cluster of the Berlin Institute of Health (BIH) using the CATALYST workflow in R^20^. Briefly, all events were clustered according to lineage marker expression using the *FlowSOM/ConsensusPlus* algorithm^21^. The number of meta-clusters was set at 30 to provide optimal granularity across the dataset. Clusters were then manually merged and annotated according to lineage marker expression and distribution of FlowSOM codes. Eighty percent of total CD45^+^ cells were neutrophils (CD66b^+^Siglec8^-^) that did not express integrin β7 (Extended Data Fig. 3f). To mitigate abundance bias, we excluded them from subsequent analyses. For visualization, dimensionality reduction was computed using the UMAP algorithm with 100 cells per sample. Cluster frequencies and marker expression were then exported for downstream analysis in R. Additionally, mass cytometry data were analyzed using FlowJo v10.4 (FlowJo).

### Cell sorting for scRNAseq

Thawed PBMCs were sorted using the SONY MA900 sorter with a 100-micron nozzle. CD45^+^ cells were sorted based on CD45 expression while eliminating DAPI-positive cells as shown in Technical Supplementary Figure 6. The sorted CD45^+^ cells were stained with the TotalSeq™ antibody mixture according to manufacturer’s protocol (BioLegend, **Suppl. Table 10**). CD4^+^ memory T cells, extracted from the same donors as those used for total PBMCs scRNAseq, were sorted on the BD FACSAria II with a 70-micron nozzle. Sorting criteria included CD3, CD4, and CD45RA expression, with the exclusion of non-viable cells by DAPI staining (Technical Supplementary Figure 6). To differentiate between individual donors, CD4^+^ memory T cells were labeled with hashtags after sorting (BioLegend, **Suppl. Table 10**).

### Single-cell sequencing of PBMCs

Following the manufacturer’s instructions, single-cell RNA-seq libraries were generated using the Chromium Next GEM Single Cell 5’ Reagent Kits v2 from 10x Genomics (Pleasanton, CA; CG000330 Rev D). Briefly, a droplet emulsion was generated in a microfluidic chip, followed by barcoded cDNA and surface tag generation within the droplets. Purified and amplified cDNA and surface tags were then subjected to NGS library construction. The manufacturer’s instructions generated T and B cell receptor sequencing libraries from amplified cDNA. Sequencing was performed on a NovaSeq 6000 instrument (Illumina). Read depth was targeted at 30K reads per cell for GEX and ∼8K for surface and TCR-BCR libraries. Clinical information regarding the patients included in the CITE-seq analysis can be found in **Suppl. Table 11**.

### scRNA-seq data analysis

Sequencing libraries for gene expression and TCR/BCR were processed together using Cell Ranger multi (v5.0.0) and the GRCh38 genome annotation and analyzed using Seurat v4.0.11^22^. We next used Seurat’s reference mapping workflow to jointly transfer cell-type labels at different granularity (“levels”) and embedding coordinates from a PBMC reference^22^ after filtering out cells with more than 10% mitochondrial gene content, less than 250 or more than 5000 genes, those with a level 1 cell type prediction score of less than 0.75, and doublets called by DoubletFinder^23^. After QC, we were left with a total of 191,578 cells for the PBMC analysis and 48,374 memory CD4^+^ T cells for downstream studies. We used scRepertoire v1.7.4 to process Cell Ranger VDJ output. ADT data were normalized using CLR normalization, and expression thresholds were determined using the Binarize R package (v1.3).

Pooled scRNAseq data for FACS-sorted CD3^+^ CD4^+^ CD45RA^-^ memory T cells from multiple donors were demultiplexed using Seurat’s HTOdemux function combined with SNP-based demultiplexing using cellSNP-lite^24^ over common variants from the 1000 Genomes Project and Vireo^25^ and otherwise processed similarly. We then selected all level 2 cell types (CD4 CTL, CD4 proliferating, CD4 TCM, CD4 TEM or Treg) from each dataset and performed a joint UMAP and clustering (at resolution 0.5) using Seurat’s SCTransform and IntegrateData workflows^26,27^. Finally, two smaller MKI67^+^ clusters were merged. Differential expression between cells in cluster 1 cells and others was performed using logistic regression, with sample type (PBMC or CD4^+^ sorted) as covariate.

### RNA isolation from tissue samples (IBDome cohort)

RNA was isolated from biopsies taken during routine endoscopy or from resected tissues at the First Department of Medicine, Friedrich-Alexander Universität Erlangen-Nürnberg (Germany), and at the Department of Gastroenterology, Infectious Diseases and Rheumatology at the Charité – Universitätsmedizin Berlin (Germany) by using a single-use biopsy forceps (Olympus). Samples were incubated in RNA protect reagent (RNAprotect Tissue Reagent, Qiagen) and stored at −80°C. One biopsy was thawed on ice and homogenized in RLT buffer (Qiagen) employing the TissueLyser LT (Qiagen) for RNA isolation. RNA was isolated using the RNeasy kit (Qiagen) and RNA Clean & Concentrator kit (Zymo Research). The concentration was measured at NanoDrop One/One (Thermo Fisher Scientific) and the quality (RNA integrity number, RIN) at Tape Station (Agilent). The RNA was used for bulk RNA sequencing at the NGS Competence Center Tübingen (NCCT).

### Bulk RNA-seq analysis

Paired sequencing reads were processed with the nf-core/rnaseq pipeline version 3.4 [10.1038/s41587-020-0439-x]. In brief, reads were aligned to the GRCh38 reference genome with GENCODE v33 annotation using STAR [10.1093/bioinformatics/bts635]. Transcripts per million (TPM) were quantified using Salmon [10.1038/nmeth.4197] and transformed to log10(TPM+1) for visualization in R.

### Proteomics assay

Proteomics analysis of patient serum samples was performed using the Olink “Target 96 Inflammation panel” platform (https://www.olink.com/products/ inflammation/). Frozen patient sera were randomly distributed on plates and spiked-in with controls according to the manufacturer’s instructions (Olink, Uppsala, Sweden). Clinical information regarding the patients included in the CITE-seq analysis can be found in **Suppl. Table 12**. The panel is a high-throughput, multiplex immunoassay that allows the simultaneous analysis of 92 inflammation-related protein biomarkers in 88 samples using Proximity Extension Assay (PEA) technology. In brief, each protein is bound by a matched pair of antibodies coupled to unique and partially complementary oligonucleotides. It hybridizes when the DNA coupled to the two antibodies is brought into close proximity. Only the tags that hybridize correctly are extended into an amplicon by quantitative real-time PCR, with a unique sequence for each protein. This dual antibody binding requirement and DNA barcoding provide exceptional readout specificity. The software then reports the relative protein concentrations of the proteins. The proteomics data is published in a normalized protein expression (NPX) format, Olink’s arbitrary unit, on the log2 scale. NPX data allows users to identify changes in individual protein levels throughout the sample and to identify protein signatures. The higher the NPX value, the higher the protein concentration. As NPX is in a log2 scale, a 1 NPX difference means a doubling in protein concentration. NPX values that did not pass internal quality control were excluded from further analysis. Downstream data analysis was performed in R using the packages limma, ggplot2 and ComplexHeatmap. The assays were done in Olink Proteomics AB, Uppsala, Sweden. Several serum proteins were different in serum when comparing IBD patients with healthy controls (HC), and were increased after vedolizumab treatment (**Technical Supplementary** Fig. 7)

### Machine learning

We aimed to identify markers across four different data modalities for predicting vedolizumab responses in patients with IBD. Therefore, we used various machine learning techniques, including both linear (logistic regression, linear support vector machines) and non-linear methods (radial SVM, polynomial SVM, random forest) and functionalities embedded in the scikit-learn-package (version 1.0.2). Logistic regression was selected as the preferred method due to its robust performance and model simplicity. Within this approach, we only considered patient data with clear response outcomes, removing patients in remission. Prior to training, incomplete patient samples were removed, and feature scaling was applied to ensure compatibility across modalities. The exact number of patients used for each run can be found in **Suppl. Table 13**. We used a 2-fold cross-validation approach for each classification run, repeated five times. Performance on the test set was evaluated using the mean area under the curve (AUC) and the standard error of the mean (SEM). Empirical p-values were calculated via permutation tests (n =1000), with AUC as the final evaluation metric. After training the classifier on the full data sets, including all parameters, we extracted the feature coefficients from the classification models. These coefficients were then used to rank and select the most influential features, comprising the top 10% of the total. A new classification model was then trained using these selected features. However, we observed that increasing the number of data modalities reduced the number of patients included due to incomplete data. To assess the robustness of the identified markers, we evaluated the classification performance using only the 13 patients with complete profiling information. The four models with the highest predictive power were identified based on the AUC. The individual feature coefficients from the reduced models were transformed into Z-scores to further evaluate the most informative markers. To validate the reliability of these final marker panels, we derived the predictive power from the flow cytometry and Olink modalities in 15 randomly selected patients. While all AUC curves were generated using the mathplotlib package (version 3.5.1), bar - and dot plots were generated using the *ggplot2* R package (version 3.3.3).

## Code availability

All original code, including mass cytometry workflows, scRNAseq analyses, and machine learning is deposited in a GitHub repository and will be publicly available upon acceptance of this manuscript.

## Data availability

All source data relevant to understanding and reproducing the results presented in this paper are provided in the supplementary data. Raw scRNAseq and mass cytometry data can be found at the following link and will be made available upon acceptance of this manuscript.

## Statistical analysis

Statistical analyses and visualizations were created in R using ggpubr, ggplot2, limma, and ComplexHeatmap packages or with the Prism Software (GraphPad Software). *P* values were calculated using the Wilcoxon test. Paired analyses are indicated by connecting lines.

## ACKNOWLEDGMENTS

We thank Nadine Sommer and Anja A. Kühl for their support with sample collection within the IBDome consortium; J. Kirsch and T. Kaiser (Flow Cytometry Core Facility, DRFZ, Berlin) for technical assistance with cell sorting; Sarah Vitcetz and Gitta Heinz from the scRNA-seq facility; Heike Hirseland and Sabine Baumgart for assistance with mass cytometry protocols, reagents, and acquisition (Mass Cytometry Group, DRFZ, Berlin); Yu-Hsien Hsieh for help with pilot experiments and analyses; and Ernesto Zarza Reyes for optimizing and revising the mass cytometry analysis code. We would like to express our gratitude to all the members of the Hegazy lab for their assistance with sample banking, as well as to our clinician and nurse colleagues in both the IBD inpatient and outpatient clinics.

A.N.H is supported by a Lichtenberg fellowship and “Corona Crisis and Beyond” grant by the Volkswagen Foundation, a BIH Clinician Scientist grant and German Research Foundation DFG-TRR241-A05 and INST 335/597-1, as well as with the ERC-StG “iMOTIONS” grant (101078069). This project was funded by a BIH Cohort Sequencing Program grant provided to A.N.H. V.H. received a Gerok fellowship from the German Research Foundation DFG-TRR241. A.L.N. was supported by the Research Training Program Stipend (Monash University). K.T. was supported by Germany’s Excellence Strategy grants EXC2151-390685813 and EXC2047-390873048. C.W. was supported by the German Research Foundation CRC-TRR241-A09, CRC-TRR241-B01 (INST 335/597-1), WE5303/3-1 as well as by the Fritz-Thyssen foundation. M.F.M. received funding from the state of Berlin and the “European Regional Development Fund” (ERDF 2014-2021, EFRE 1.8/11, Deutsches Rheuma-Forschungszentrum); the Leibniz Association (Leibniz Collaborative Excellence, TargArt and ImpACt); and the German Federal Ministry of Education and Research (BMBF) projects CONAN and TReAT. B.S. was supported by the German Research Foundation CRC-TRR241 (project-ID: 375876048), CRC1340 (project-ID: 372486779), CRC1449 (project-ID: 321232613), and INST 335/597-1. BS received Lecture Fees from Abbvie, BMS, CED Service GmbH, Chiesi, Falk, Forga Software, Galapagos, IBD Passport, Janssen, Materia Prima, Pfizer, and Lilly; Consultancy fees: Abbvie, Arena Pharma, Boehringer, BMS, Celgene, CT-Scout, Endpoint Health, Galpagos, Gilead, Janssen, Landos, Lilly, Pfizer, PredictImmune, and PsiCro, and research support from Pfizer (all payments went to institution). The authors declare that no competing interests exist.

## AUTHOR CONTRIBUTIONS

Study design and conceptualization: ANH, VH; writing—original draft preparation: VH, ANH with input from CC, LS, BO, KT; commenting—and editing: LS, CW, HM, ZT, BS, KT; consenting patients, providing and interpreting clinical data: KF, VH, DL, ES, CW, BS, ALN, ANH; visualization: ANH, VH, CC, LS, BW, CP; scRNAseq library and sequencing: TC, MFM; scRNAseq bioinformatic analysis: BO; Bulk RNAseq analysis: CP, ZT; CyTOF acquisition and data analysis: VH; CyTOF protocols, acquisition, and reagents: VH, MB, AS, HEM; data integration and machine learning: LS, KT; FACS staining, acquisition and analysis: CC, KF; sample biobanking: KF, VH, DB, EM, ES, ALN. All authors revised the manuscript and approved the final version submitted for publication.

## OTHER CONTRIBUTING AUTHORS

TRR241 IBDome Consortium: Imke Atreya^1^, Raja Atreya^1^, Petra Bacher^2,3^, Christoph Becker^1^, Christian Bojarski^4^, Nathalie Britzen-Laurent^1^, Caroline Bosch-Voskens^1^, Hyun-Dong Chang^5^, Andreas Diefenbach^6^, Claudia Günther^1^, Ahmed N. Hegazy^4^, Kai Hildner^1^, Christoph S. N. Klose^6^, Kristina Koop^1^, Susanne Krug^4^, Anja A. Kühl^7^, Moritz Leppkes^1^, Rocío López-Posadas^1^, Leif S.-H. Ludwig^8,9^, Clemens Neufert^1^, Markus Neurath^1^, Jay Patankar^1^, Magdalena Prüß^3^, Andreas Radbruch^5^, Chiara Romagnani^3^, Francesca Ronchi^6^, Ashley Sanders^4,9^, Alexander Scheffold^2^, Jörg-Dieter Schulzke^4^, Michael Schumann^4^, Sebastian Schürmann^1^, Britta Siegmund^4^, Michael Stürzl^1^, Zlatko Trajanoski^10^, Antigoni Triantafyllopoulou^5,11^, Maximilian Waldner^1^, Carl Weidinger^4^, Stefan Wirtz^1^, Sebastian Zundler^1^

^1^Department of Medicine 1, Friedrich-Alexander University, Erlangen, Germany

^2^Institute of Clinical Molecular Biology, Christian-Albrecht University of Kiel, Kiel, Germany. ^3^Institute of Immunology, Christian-Albrecht University of Kiel and UKSH Schleswig-Holstein, Kiel, Germany

^4^Charité – Universitätsmedizin Berlin, corporate member of Freie Universität Berlin and Humboldt-Universität zu Berlin, Department of Gastroenterology, Infectious Diseases and Rheumatology, Berlin, Germany

^5^Deutsches Rheuma-Forschungszentrum, ein Institut der Leibniz-Gemeinschaft, Berlin, Germany

^6^Charité – Universitätsmedizin Berlin, corporate member of Freie Universität Berlin and Humboldt-Universität zu Berlin, Institute of Microbiology, Infectious Diseases and Immunology, Berlin, Germany

^7^Charité - Universitätsmedizin Berlin, corporate member of Freie Universität Berlin and Humboldt-Universität zu Berlin, iPATH.Berlin, Berlin, Germany

^8^Berlin Institute of Health at Charité-Universitätsmedizin Berlin, Berlin, Germany.

^9^Max Delbrück Center for Molecular Medicine in the Helmholtz Association (MDC), Berlin Institute for Medical Systems Biology (BIMSB), Berlin, Germany

^10^Biocenter, Institute of Bioinformatics, Medical University of Innsbruck, Innsbruck, Austria. ^11^Charité – Universitätsmedizin Berlin, corporate member of Freie Universität Berlin and Humboldt-Universität zu Berlin, Department of Rheumatology and Clinical Immunology, Berlin, Germany

