## Supplementary figures for "Multimodal profiling of peripheral blood identifies proliferating circulating effector CD4^+^ T cells as predictors for response to integrin α4β7-blocking therapy in patients with inflammatory bowel disease"

### EXTENDED DATA FIGURES 1-10 AND TECHNICAL SUPPLEMENTARY FIGURES 1-8

#### Multimodal profiling of peripheral blood identifies proliferating circulating effector CD4$^+$ T cells as predictors for response to integrin α4β7-blocking therapy in patients with inflammatory bowel disease

Veronika Horn[1,2,13,14], Camila Cancino[1,2,13], Lisa Steinheuer[3,13], Benedikt Obermayer[4,13], *et al*

[1] Charité – Universitätsmedizin Berlin, corporate member of Freie Universität Berlin and Humboldt-Universität zu Berlin, Department of Gastroenterology, Infectious Diseases and Rheumatology, 12203 Berlin, Germany

[2] Deutsches Rheuma-Forschungszentrum, ein Institut der Leibniz-Gemeinschaft, 10117 Berlin, Germany

[3] Institute for Experimental Oncology, Biomathematics Division, University Hospital Bonn, Bonn, Germany

[4] Berlin Institute of Health at Charité – Universitätsmedizin Berlin, Core Unit Bioinformatics, 10117 Berlin, Germany

[13] Equal contribution

**Corresponding author:** Prof. Ahmed N. Hegazy, M.D, Ph.D., Department of Gastroenterology, Infectious Diseases and Rheumatology, Charité - Universitätsmedizin Berlin, Campus Benjamin Franklin, Hindenburgdamm 30, 12203 Berlin, Germany; and Deutsches Rheuma-Forschungszentrum, ein Institut der Leibniz-Gemeinschaft, 10117 Berlin, Germany. Phone: +49 30 450 614 378;

**EXTENDED DATA FIGURES 1-10**

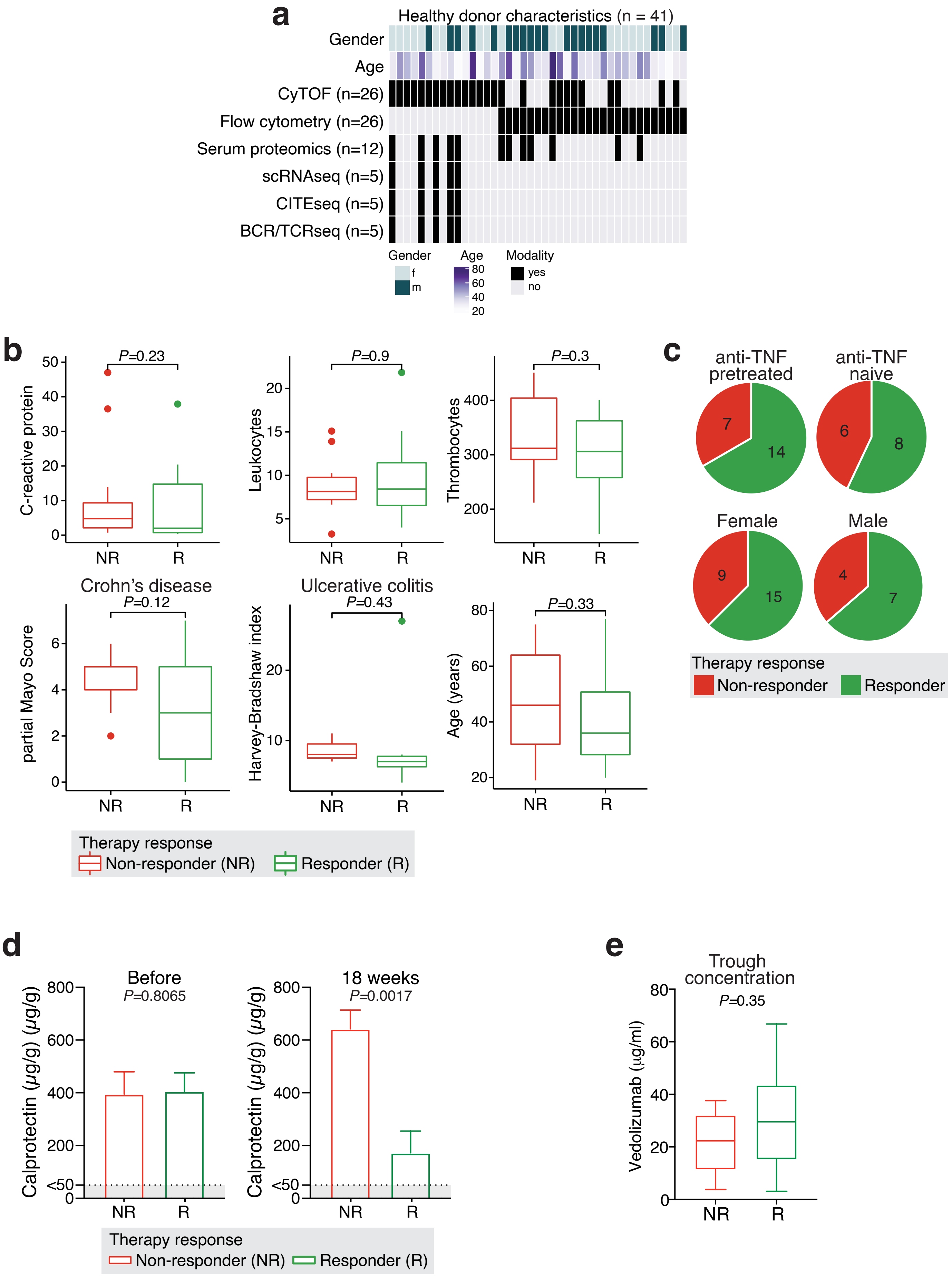


Extended Data Fig. 1

**Extended Data Figure 1. Characteristics of healthy donors and (para-)clinical data of IBD patients.**

(a) Graph depicting the age and gender distribution within the healthy control cohort (n=41), along with the techniques applied to each sample.

(b) Box plots illustrating the comparison of (para-)clinical parameters between non-responders and responders prior to therapy (responders n=22, non-responders n=13).

(c) Pie charts illustrating the indicated characteristics in IBD patients before therapy.

(d) Bar graph illustrating stool calprotectin levels before initiation of treatment and approximately 18 weeks after treatment commencement.

(e) Trough concentrations in IBD patients receiving vedolizumab at week 6 of treatment. (b, d) Mann-Whitney test.

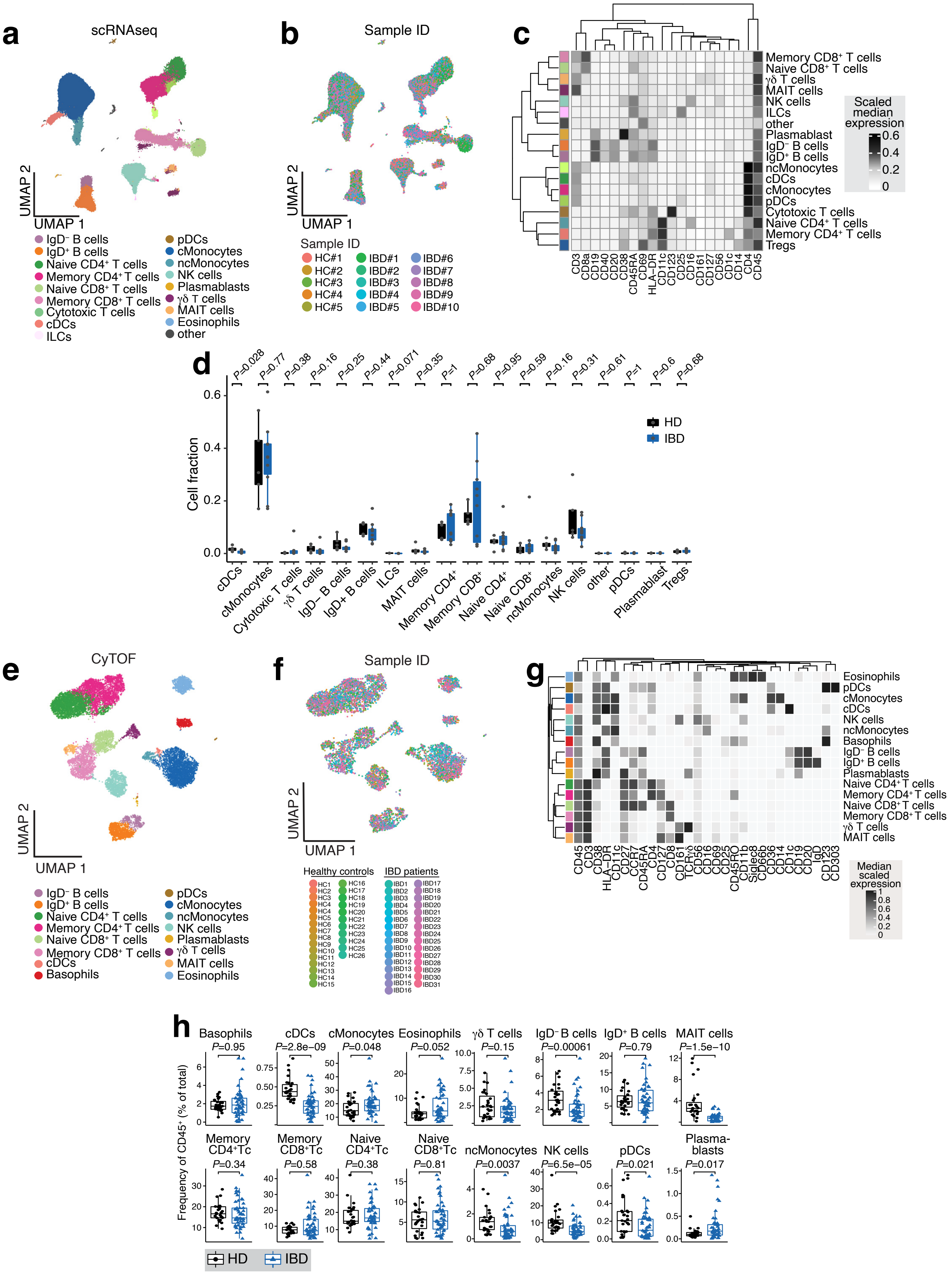


Extended Data Fig. 2

**Extended Data Fig. 2. Immune cell subset identification and quantification in peripheral blood of healthy controls and IBD patients.**

(a, b) UMAP plots of CITE-seq profiled peripheral blood cells with color-coded representation of cell types and patients (n=191,578 cells derived from a total of 25 samples: HC, n=5; IBD, n=10 matched before and after treatment). Sorted $CD45^+$ cells from the peripheral blood of 5 healthy controls and 10 ulcerative colitis patients before and 6 weeks after vedolizumab (VDZ) treatment initiation were subjected to CITE-seq analysis.

(c) Heatmap of surface antigen expression on identified cell types in peripheral blood cells derived from CITE-seq data.

(d) Box plot of cell type proportions comparing healthy controls and IBD patients. Mann Whitney test.

(e, f) UMAP plots of peripheral blood cells extracted from the CyTOF dataset, with color representation based on cell type and patient. Results derived from FlowSOM/ConsensusPlus clustering analysis performed on 13,311,287 cells from 154 samples. This dataset includes both healthy controls and IBD patients both before and during treatment.

(g) Median scaled lineage marker expression in the indicated cluster resulting from FlowSOM clustering.

(h) Comparison of cluster frequencies between healthy control (n=27) and IBD patients (n=31) before therapy. Mann Whitney test.

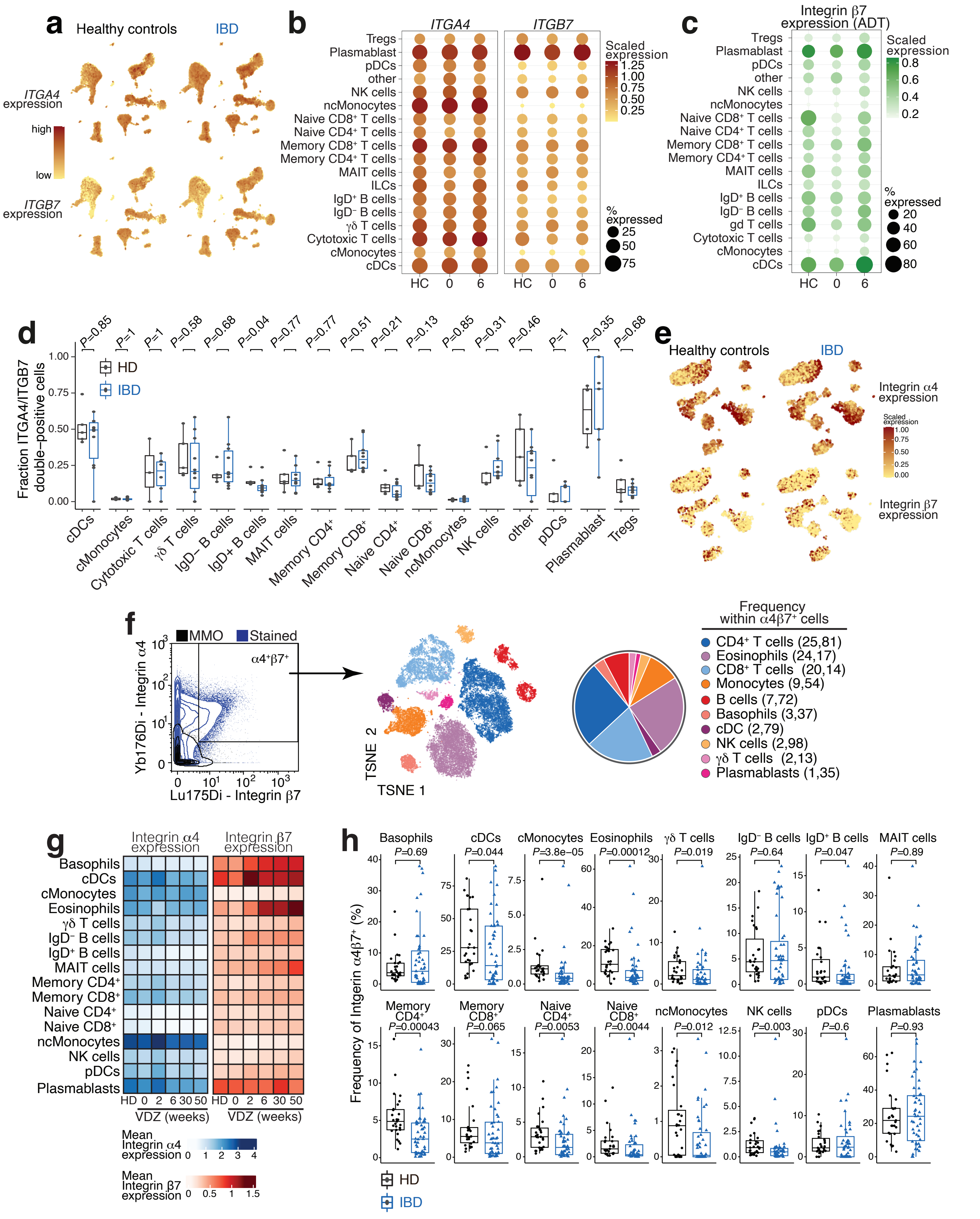


Extended Data Fig. 3

**Extended Data Fig. 3. Integrin α4β7 expression and distribution in circulating immune cells at steady state and after vedolizumab treatment.**

(a) Expression of *ITGA4* and *ITGB7* in the different cell subsets derived from scRNA-seq data.

(b) Dot plot of *ITGA4* and *ITGB7* expression in scRNA-seq data from different cell types and different conditions; color indicates expression level, dot size the percentage of positive cells.

(c) As in (b), derived from integrin β7 protein expression (antibody-derived tag, ADT) by CITE-seq.

(d) Box plot of the proportion of *ITGA4/ITGB7* double-positive cells in scRNA-seq data, comparing healthy controls and IBD patients. Mann-Whitney test.

(e) Scaled expression of integrin α4 and β7 in the CyTOF dataset from healthy control and IBD patients before therapy (IBD).

(f) Representative dot plot of integrin α4 (CD49d) and β7 staining in mass cytometry on $CD45^+$ cells showing stained cells (blue) and metals-minus-one (MMO) control. TSNE with results of FlowSOM clustering of $\alpha4\beta7^+$ cells from the mass cytometry dataset and a pie chart representing the percentage of cluster frequency within the total $\alpha4\beta7^+$ cells derived from HC and IBD patients.

(g) Mean integrin α4 and β7 expression within mass cytometry clusters, as determined by the FlowSOM algorithm in healthy controls and IBD patients before treatment and at the indicated time points after treatment initiation (HC=27, before treatment n=31, week 2 n=9, week 6 n=25, week 30 n=15, week 50 n=9).

(h) Frequency of integrin α4β7 double-positive cells from the indicated cell populations in mass cytometry data (HC, n=27; IBD, n=31). Mann-Whitney test.

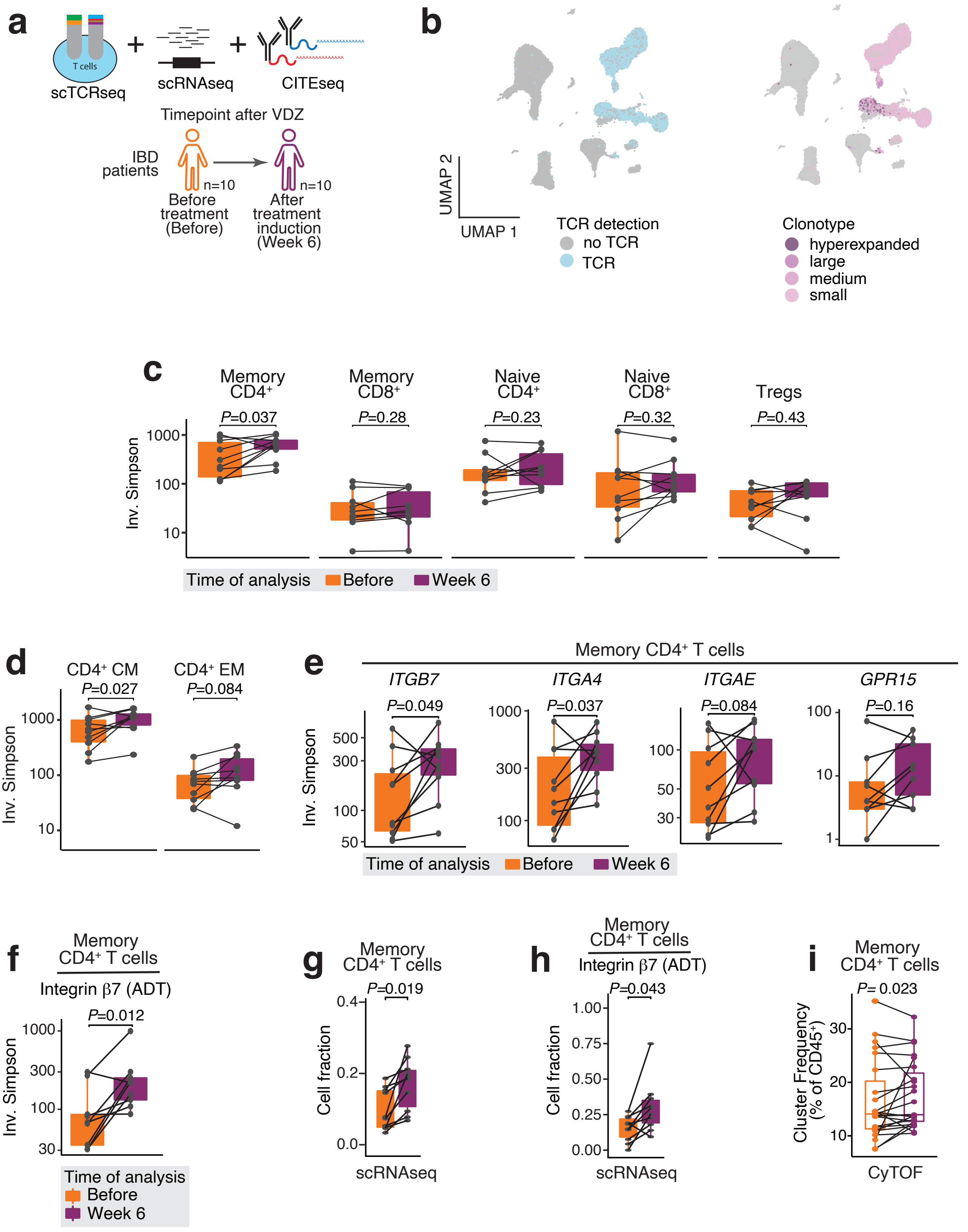


Extended Data Fig. 4

**Extended Data Fig. 4. Vedolizumab enhances clonal diversity among circulating $CD4^+$ memory T cells.**

(a) Schematic of the experimental approach.

(b) UMAP plot indicating cells with detected TCR sequence (left) and frequency of the associated clonotype (right; small: $< 10^{-3}$, medium: $< 10^{-2}$, large: $< 10^{-1}$, hyperexpanded: $> 10^{-1}$).

(c) Clonal diversity quantified by the inverse Simpson index in various T cell subsets before treatment and 6 weeks after treatment.

(d) Clonal diversity in central and effector memory $CD4^+$ T cell subpopulations.

(e) Clonal diversity in memory $CD4^+$ T cell subpopulations defined by expression of the indicated marker genes.

(f) Clonal diversity in memory $CD4^+$ T cells with positive integrin β7 expression (ADT) determined by CITE-seq.

(g) Abundance of memory $CD4^+$ T cells among all sequenced cells per sample.

(h) Abundance of memory $CD4^+$ T cells with positive integrin β7 expression determined by CITE-seq.

(i) Cluster frequency in the mass cytometry dataset.

(c-i) Wilcoxon test.

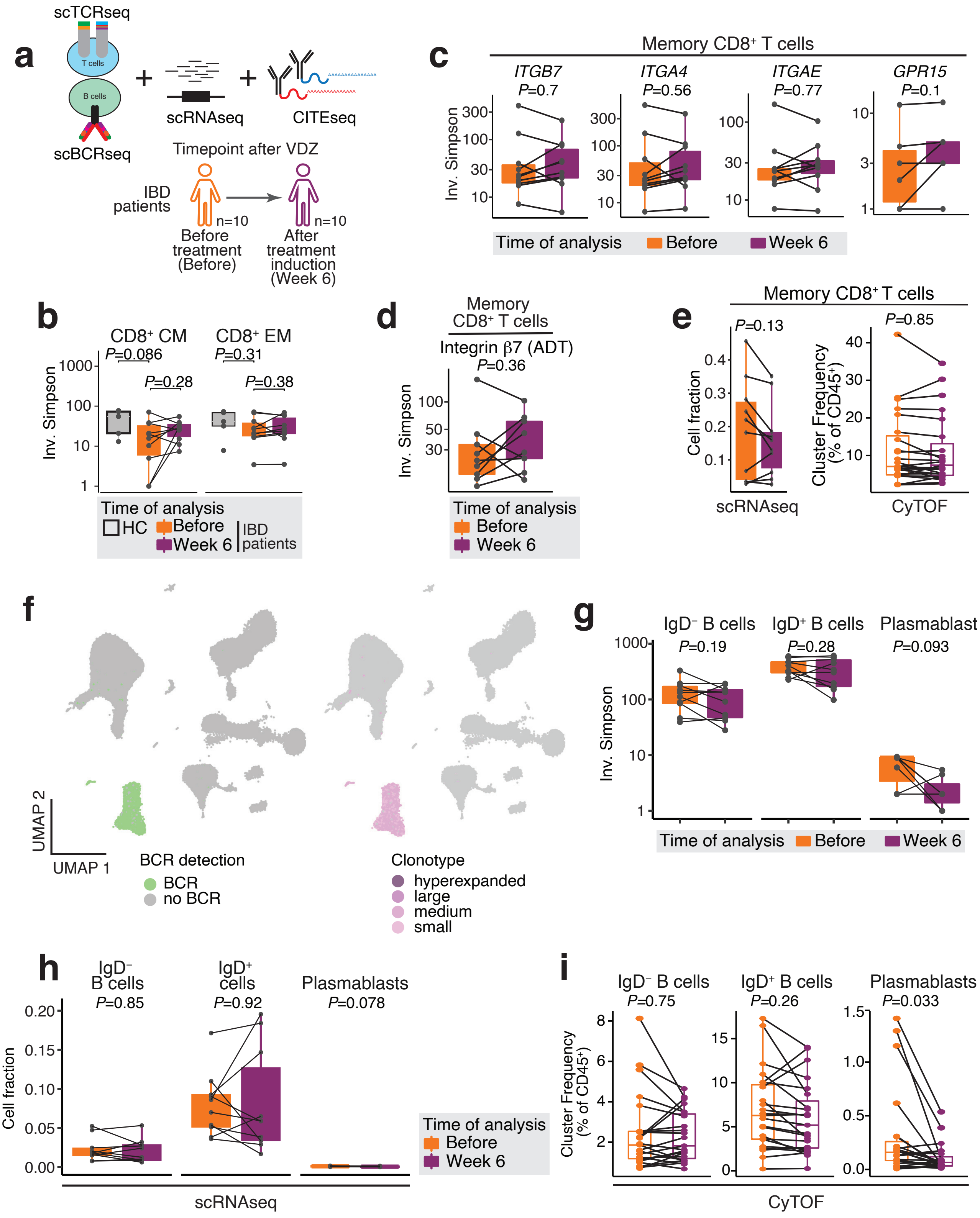


Extended Data Fig. 5

**Extended Data Fig. 5. Vedolizumab treatment does not affect the diversity of circulating CD8 T or B cells.**

(a) Schematic of the experimental approach.

(b) Clonal diversity quantified by the inverse Simpson index in central and effector memory $CD8^+$ T cell subpopulations.

(c) Box plot of TCR repertoire diversity in memory $CD8^+$ T cells positive for the indicated markers.

(d) Box plot of TCR repertoire diversity in memory $CD8^+$ T cells positive for integrin β7 (ADT) determined by CITE-seq.

(e) Box plot depicting the abundance of $CD8^+$ memory T cells before and 6 weeks after vedolizumab treatment using both scRNA-seq data (left) and CyTOF data (right).

(f) UMAP plots of scRNA-seq data showing cells with detected BCR sequence (left) and expansion status of the associated clonotype (right; small: $< 10^{-3}$, medium: $< 10^{-2}$, large: $< 10^{-1}$, hyperexpanded: $> 10^{-1}$).

(g) Box plot of BCR repertoire diversity quantified by the inverse Simpson index in the indicated B cell subpopulations before and 6 weeks after starting vedolizumab treatment.

(h) Box plot of the proportions of the indicated B cell types before and after vedolizumab treatment.

(i) Box plot depicting the abundance of B cell subtypes before and 6 weeks after vedolizumab treatment using CyTOF data.

(b-e, g-i) Wilcoxon test.

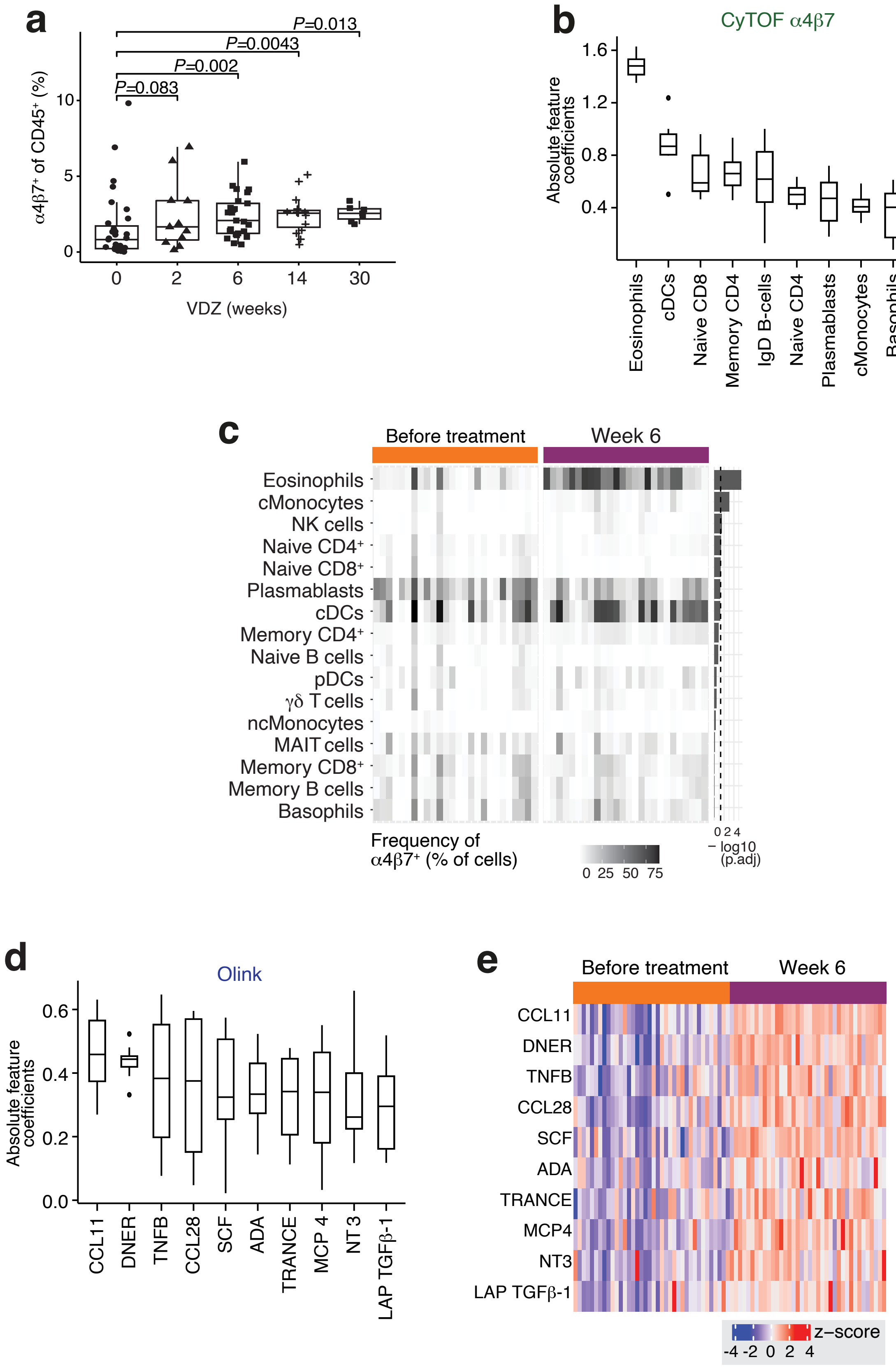


Extended Data Fig. 6

**Extended Data Fig. 6. Vedolizumab modulates the abundance of circulating integrin α4β7-positive cells and serum proinflammatory proteins.**

(a) Percentage of $\alpha4\beta7^{+}$ cells from total $CD45^{+}$ cells in the mass cytometry dataset at different timepoints throughout treatment (IBD patients before treatment n=31, week 2 n=9, week 6 n=25, week 30 n=15, week 50 n=9). Mann-Whitney test.

(b) Overview of the top ten cell populations with the most significant changes in integrin α4β7 expression identified from the CyTOF data. These populations have the highest feature coefficients, and the absolute values of these coefficients are shown. The complete list can be found in **Suppl. Table 2**.

(c) Heatmap showing the abundance of integrin $\alpha4\beta7^{+}$ cells within the indicated cell populations (IBD patients before treatment n=31; after treatment n=25). Wilcoxon test with FDR correction.

(d) Overview of the ten vedolizumab efficacy features from Olink data with the largest feature coefficients.

(e) Olink quantification of the indicated analytes before treatment (n=39) and 6 weeks after vedolizumab treatment (n=39).

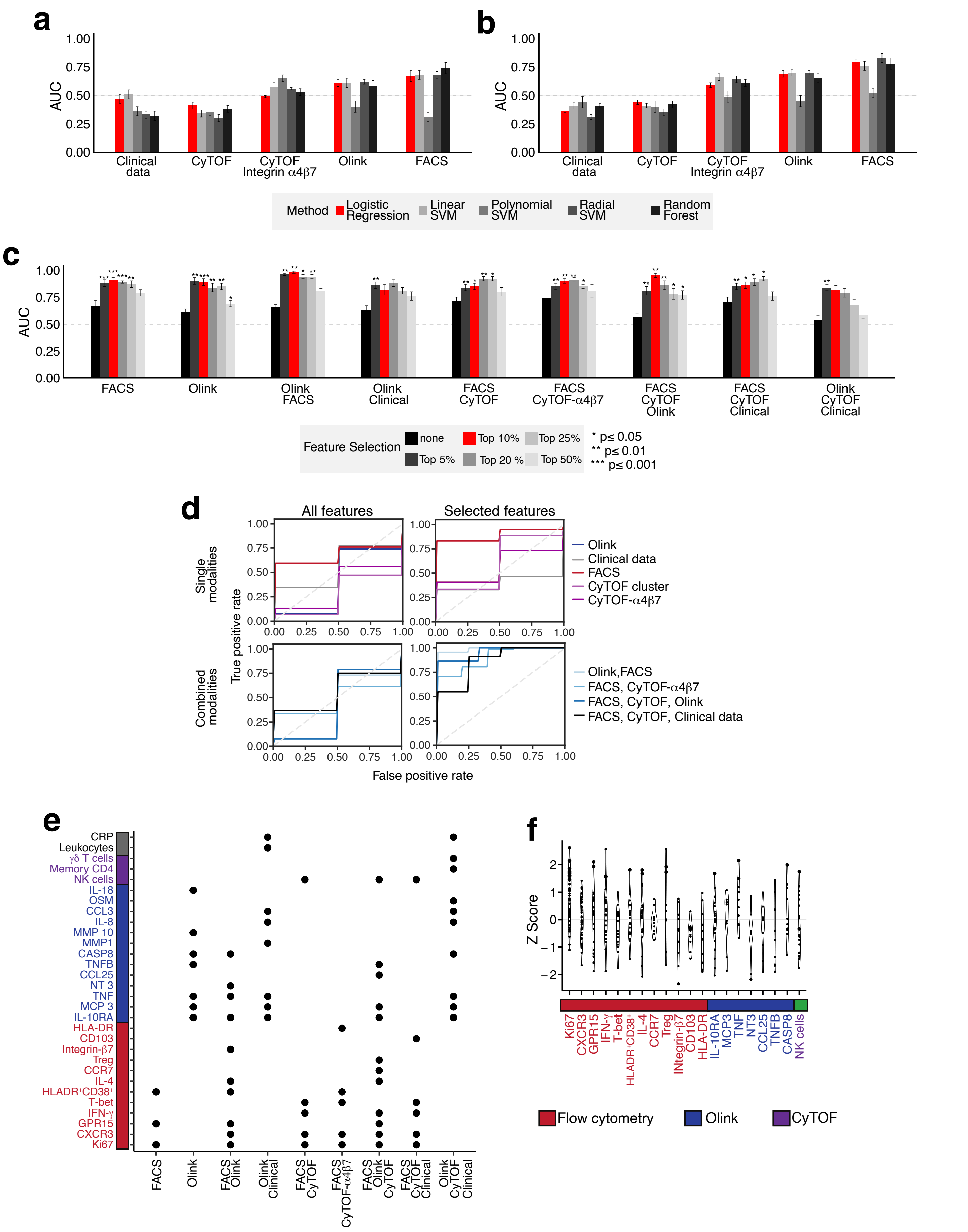


Extended Data Fig. 7

**Extended Data Fig. 7. Technical approach showing data integration and optimization of classification tools.**

(a) Comparison of five different classification tools within individual data sets. Shown are AUC values with SEM.

(b) Comparison of five classification tools using selected features. Shown are AUC values with SEM. Feature importance was determined using logistic regression, and only those in the top 50% were retained. The color code is the same as in panel A. Due to similar mean AUC scores across data sets and the simplicity of the model, logistic regression (red bars) was selected as the preferred classification tool in subsequent analyses.

(c) Predictive ability before and after various feature engineering thresholds. Per model, all features were ranked according to their absolute feature coefficients. Models that did not have an AUC of at least 0.5 (average AUC minus SEM) as indicated by the dashed line are not shown. Empirical *p*-values were calculated using permutation tests (n=1000). Based on the robustness of the obtained AUCs, we used the top 10% cutoff (red bars).

(d) Predictive capacity of the models considering completely profiled patients (n=13), analogous to Fig. 1e.

(e) Overview of most influential features in the classification models shown in (b). For each data set, a feature is selected if its absolute feature coefficient scored within the top 10% of all features.

(f) Scatterplot of the Z-score transformed feature coefficients shown in **Fig. 1f**.

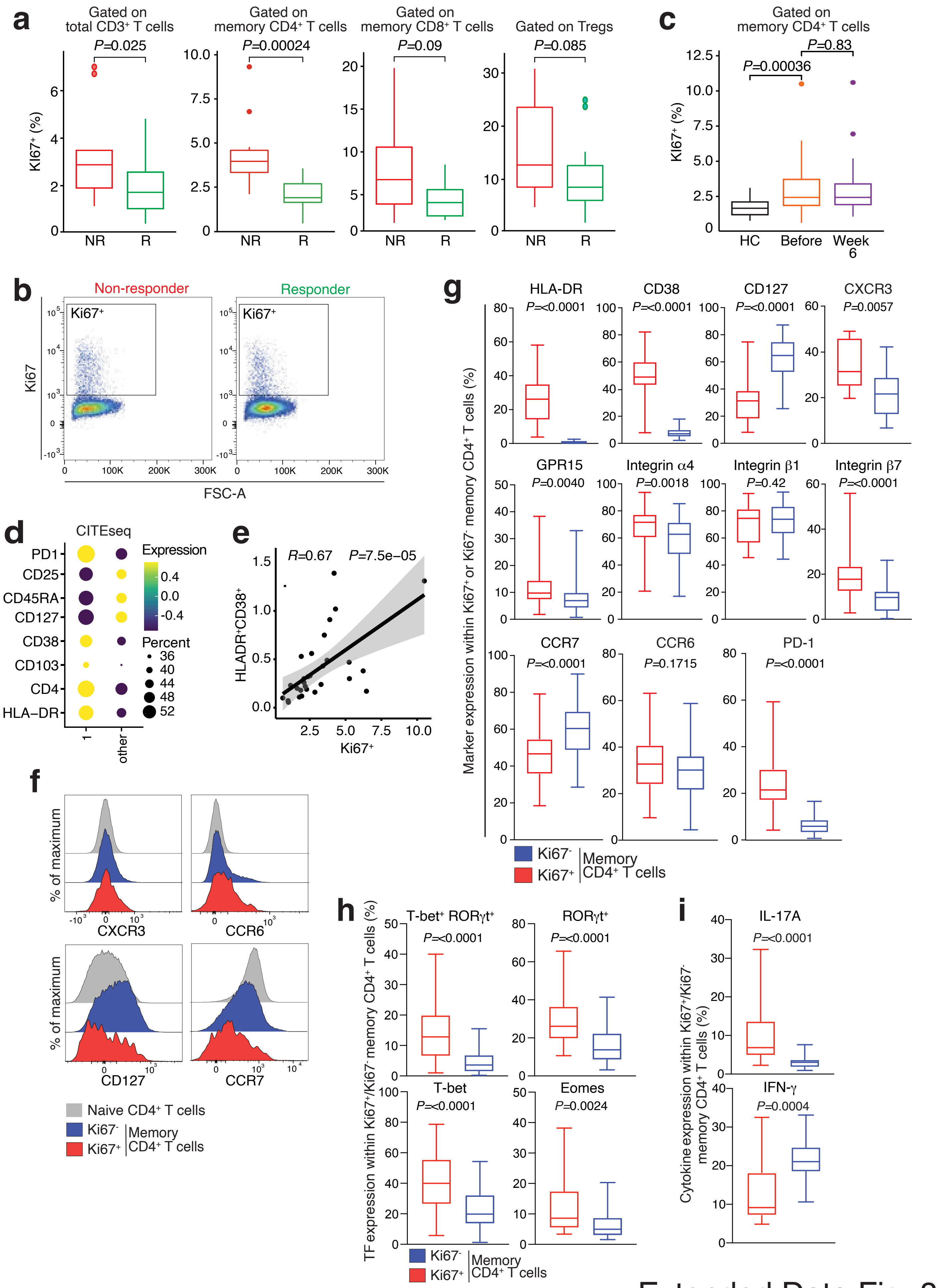


Extended Data Fig. 8

**Extended Data Figure 8. Distinctive features of proliferating effector $CD4^+$ T cells in vedolizumab non-responsive patients.**

(a) Box plots illustrating the percentage of $Ki67^+$ cells within the indicated T cell subsets among IBD patients prior to initiating therapy (responders n=19, non-responders n=10).

(b) FACS plot showing Ki67 expression in memory $CD4^+$ T cells in representative donors.

(c) Box plots depicting the percentage of $Ki67^+$ cells within memory $CD4^+$ T cells across three groups: healthy controls (HC, n=26), IBD patients before treatment (n=41), and IBD patients 6 weeks after treatment initiation (n=35).

(d) Dot plot displaying scaled CITE-seq expression of selected markers in cluster 1 in comparison to other clusters.

(e) Scatter plot depicting the correlation between the percentage of $HLA\text{-}DR^+CD38^+$ cells within memory $CD4^+$ T cells and the frequency of Ki67-positive cells within memory CD4 T cells of IBD patients (n=41). This correlation was evaluated using Spearman's rank correlation analysis based on data obtained by flow cytometry.

(f) Histograms showcasing FACS surface marker expression in a representative IBD patient. The plot displays the expression of specified markers within naïve, $Ki67^+$, and $Ki67^-$ memory $CD4^+$ T cells.

(g) Percentage of indicated surface marker expression within $Ki67^+$ and $Ki67^-$ memory $CD4^+$ T cells in IBD patients. n=29-44. Mann-Whitney test.

(h) Percentage of indicated transcription factor expression within $Ki67^+$ and $Ki67^-$ memory $CD4^+$ T cells in IBD patients. n=28-40.

(i) Percentage of IL-17A and IFN-γ expression within $Ki67^+$ and $Ki67^-$ memory $CD4^+$ T cells in IBD patients. n=20.

(a, c, g, h) The statistical significance was assessed using the Mann-Whitney test.

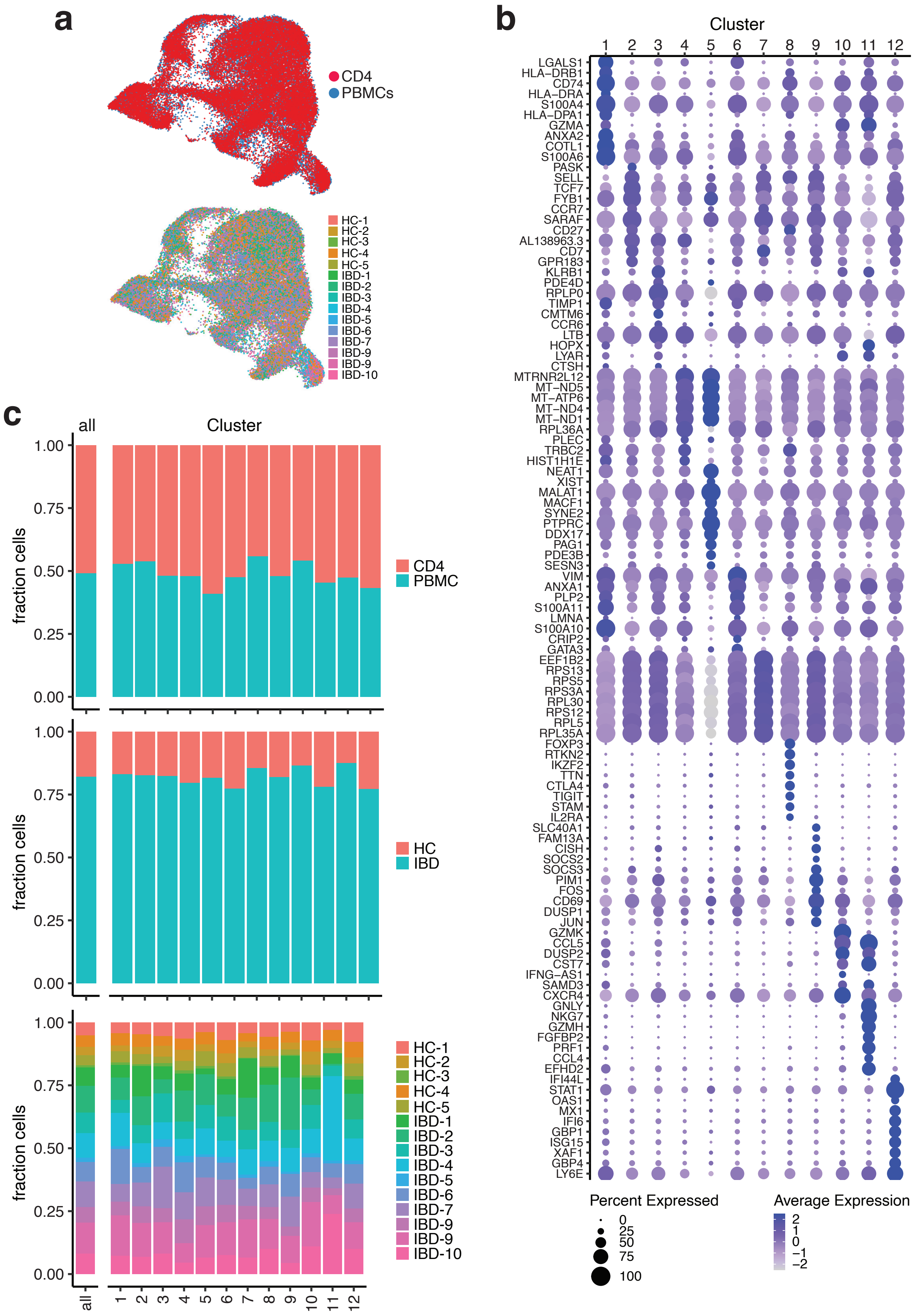


Extended Data Fig. 9

**Extended Data Figure 9. Characterization of $CD4^+$ memory T cells through single-cell RNA sequencing (scRNA-seq).**

(a) UMAP plots of scRNA-seq profiles of memory $CD4^+$ T cells derived from total $CD45^+$ or sorted $CD3^+CD4^+CD45RA^{low}$ cells with color-coded representation of cell source (CD or PBMC) and patients (data derived from 25 samples: HC, n=5; IBD, n=10 matched before and after treatment). Sorted $CD45^+$ and $CD3^+CD4^+CD45RA^{low}$ cells from matched peripheral blood of 5 healthy controls and 10 ulcerative colitis patients before and 6 weeks after vedolizumab (VDZ) treatment initiation were subjected to scRNA-seq analysis.

(b) Dot plot of cluster marker genes for the $CD4^+$-memory T cell subset in scRNA-seq data from PBMCs and memory $CD4^+$-sorted cells. Dot size indicates percentage of positive cells, color scale indicates scaled average expression.

(c) Proportions of cells derived from PBMCs or $CD4^+$ memory T cells in healthy versus IBD patients and different donors.

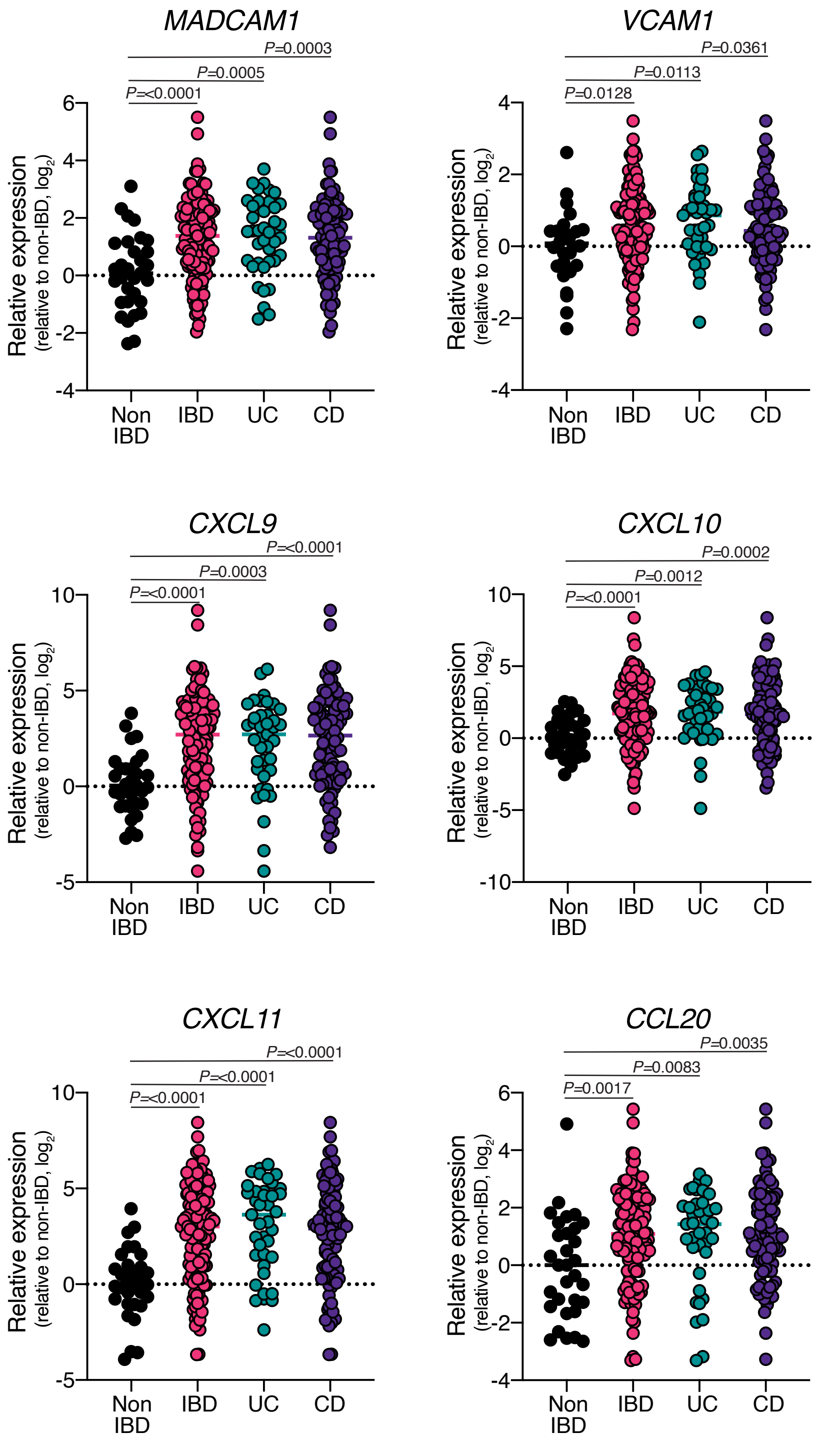


Extended Data Fig. 10

**Extended Data Figure 10. Expression of chemokine ligands and adhesion molecules in mucosal tissues of non-IBD controls and IBD patients.**

Expression of the indicated genes in mucosal samples obtained from non-IBD controls (n=32), UC patients (n=39), and CD patients (n=85). Endoscopic biopsies and surgical specimens were collected within the IBDome cohort and subjected to RNA sequencing. The gene expression in UC and CD samples were normalized to the median expression level in non-IBD control samples. Statistical comparisons were made using one-way ANOVA with Dunn's multiple-comparison tests.

**TECHNICAL SUPPLEMENTARY FIGURES 1-8**

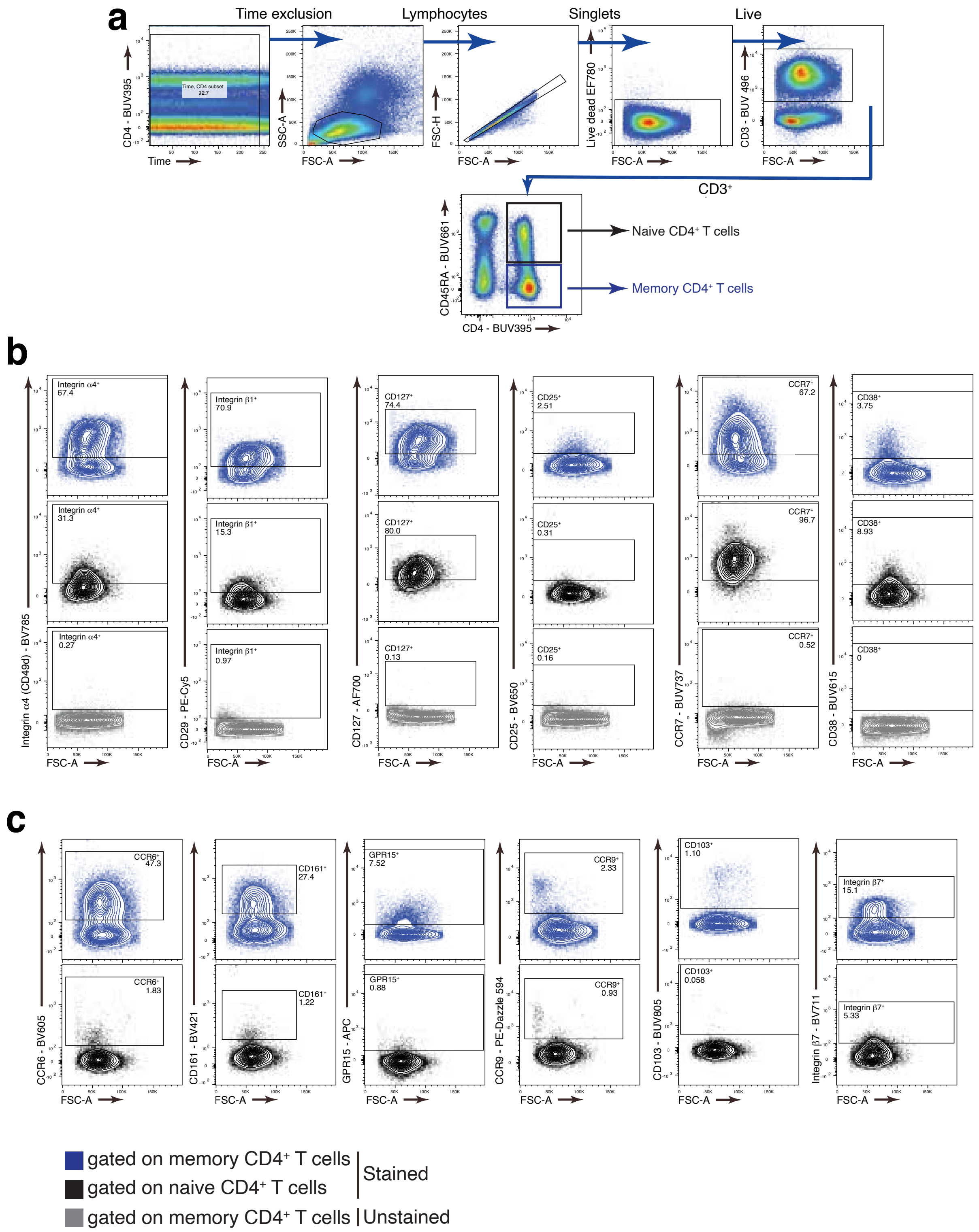


### Technical Supplementary Figure 1

**Technical Supplementary Fig. 1. Identification of peripheral blood CD4$^+$ memory T cells via flow cytometry.**

(a) Gating strategy for CD4$^+$ memory T cells shown for one representative donor.

(b, c) Dot plots depicting the expression of the indicated surface markers. Naïve, memory CD4$^+$ T cells, and staining control are shown for each surface marker for one representative donor.

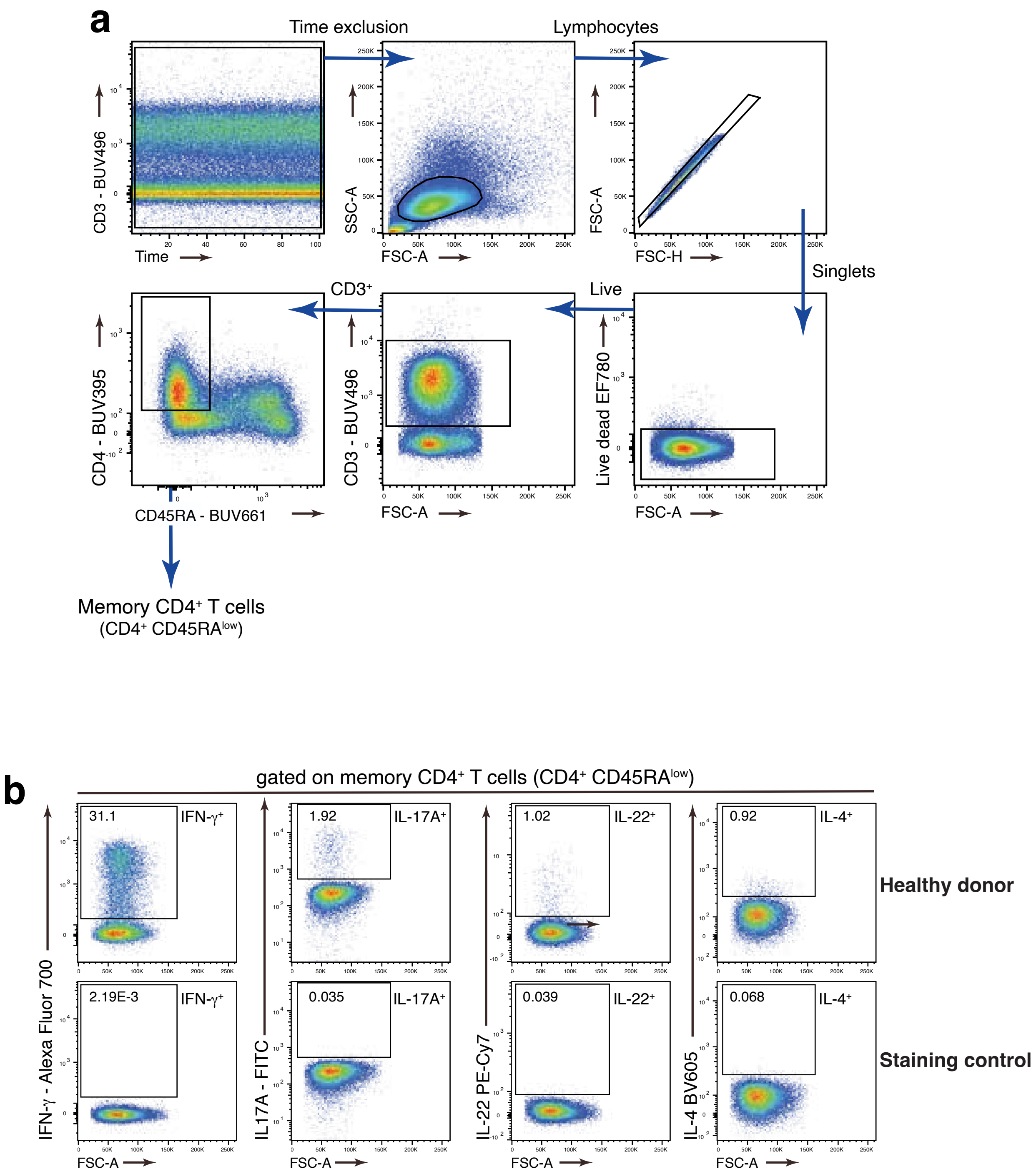


Technical Supplementary Figure 2

**Technical Supplementary Fig. 2. Exploring cytokine expression in $CD4^+$ memory T cells using intracellular cytokine staining and flow cytometry.**

(a) Gating strategy for $CD4^+$ memory T cells after stimulation with PMA/ionomycin and intracellular staining.

(b) Dot plots depicting the expression of the indicated cytokines. $CD4^+$ memory T cells, and staining control (no stimulation control) are shown for each cytokine.

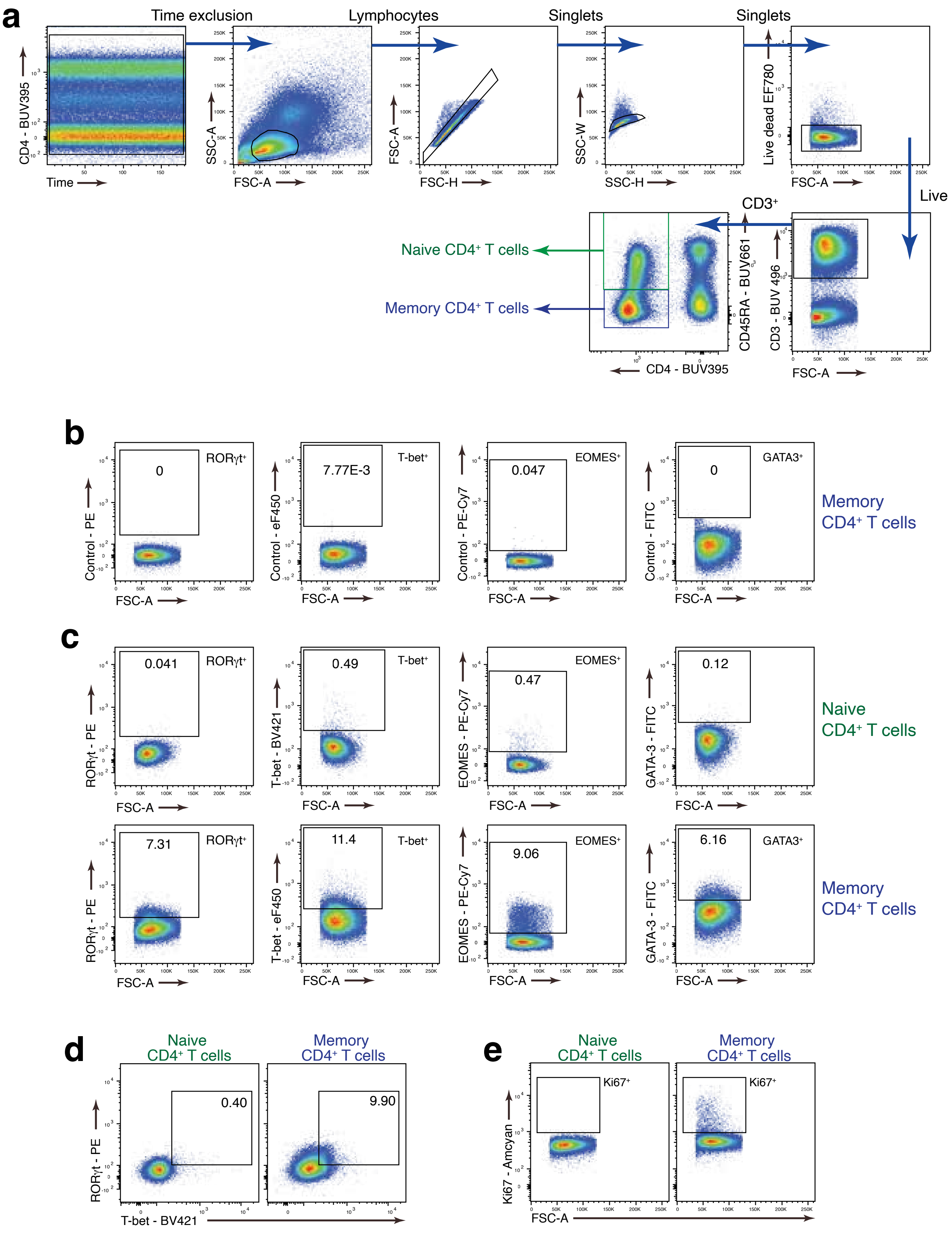


Technical Supplementary Figure 3

**Technical Supplementary Fig. 3. Evaluating the expression of transcription factors in $CD4^+$ memory T cells using flow cytometry.**

(a) Gating strategy for memory $CD4^+$ T cells after fixation with eBioscience™ Foxp3/Transcription Factor Staining Buffer Set and intracellular staining.

(b) Cells were stained with surface markers, and the indicated channels for the transcription factors were left empty. Gated on $CD4^+$ memory T cells.

(c) Dot plots depicting the expression of the indicated transcription factors. Cells were stained for the indicated transcription factors. Shown are naïve and memory $CD4^+$ T cells.

(d) Dot plots showing the co-expression of RORγt and T-bet in naïve and memory $CD4^+$ T cells.

(e) Dot plots showing the expression of Ki67 in naïve and memory $CD4^+$ T cells.

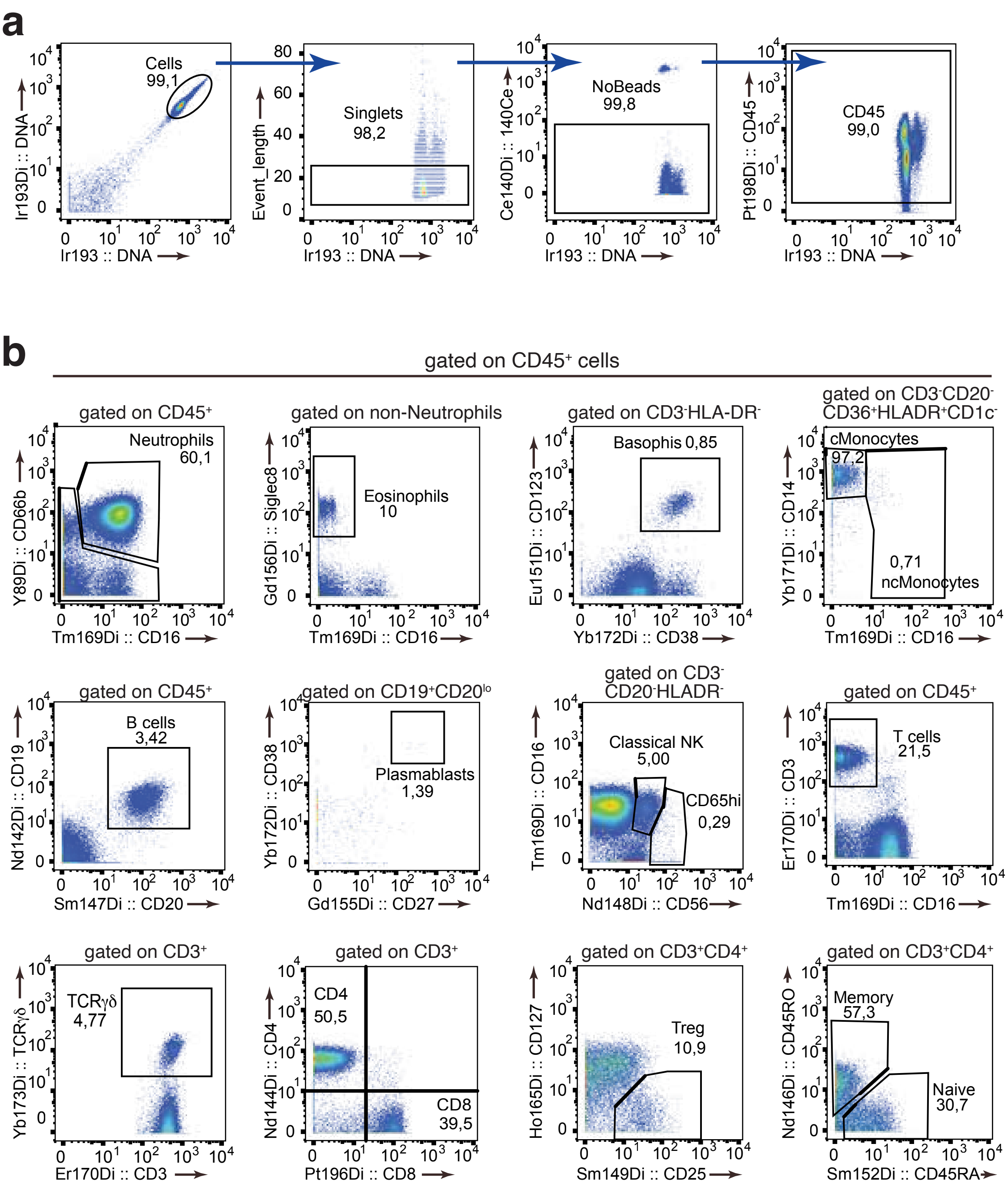


Technical Supplementary Figure 4

**Technical Supplementary Fig. 4. Immune cell subtype identification using mass cytometry.**

(a) Gating approach and identification of $CD45^+$ cells.

(b) Dot plots showing the expression of the indicated markers and the identification of the depicted cell populations.

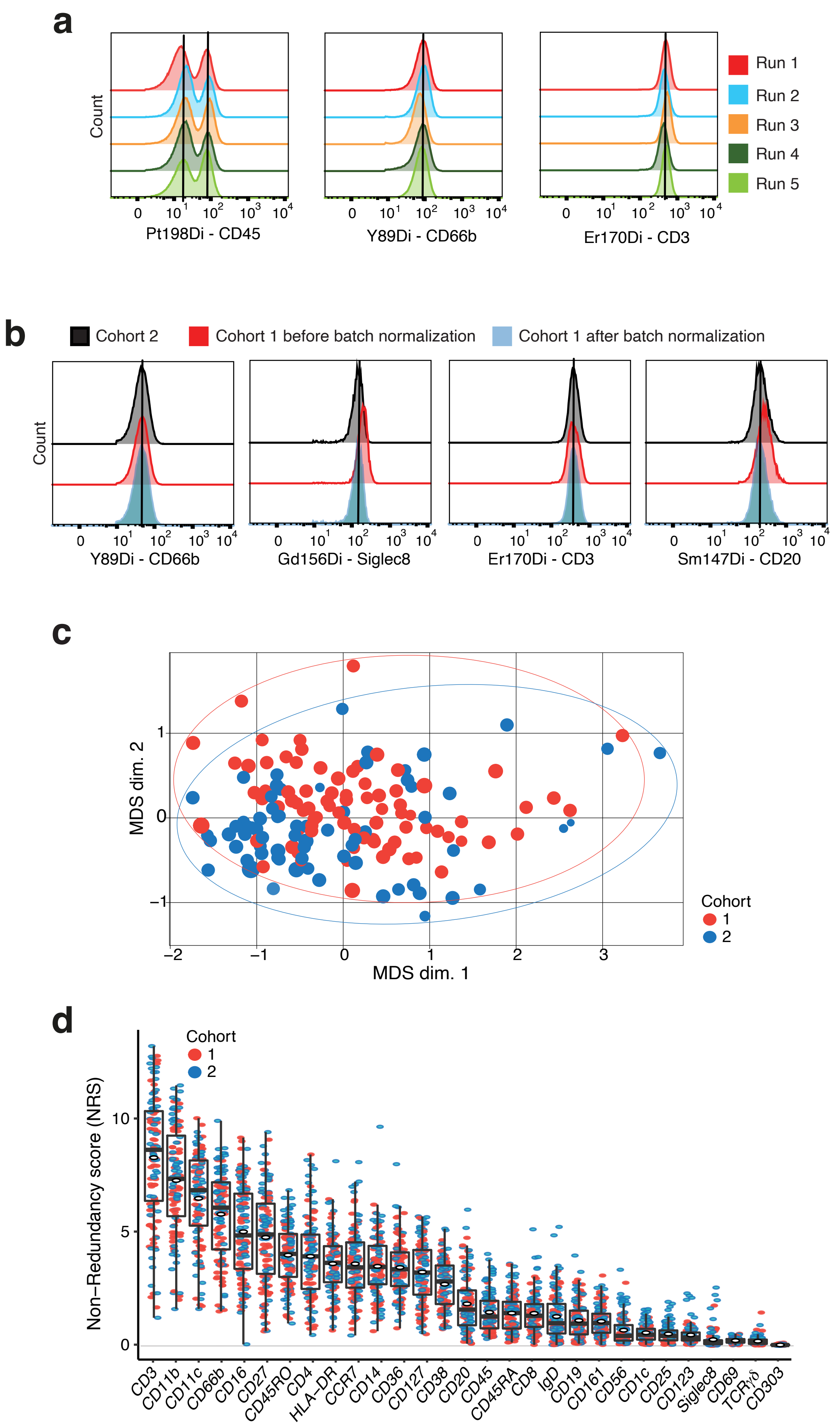


Technical Supplementary Figure 5

**Technical Supplementary Fig. 5. Quality control for mass cytometry data acquisition.**

(a) Histograms depicting expression of exemplary lineage markers among anchor samples (from the same donor) across different runs within a single cohort.

(b) Histograms depicting expression of exemplary lineage markers within the same donor across different cohorts, both before and after batch normalization.

(c) MDS plot showing the distribution of both cohorts after batch normalization. Each sample represents one plot.

(d) Boxplot of lineage marker distribution calculated by non-redundancy score (NRS) between cohorts 1 and 2.

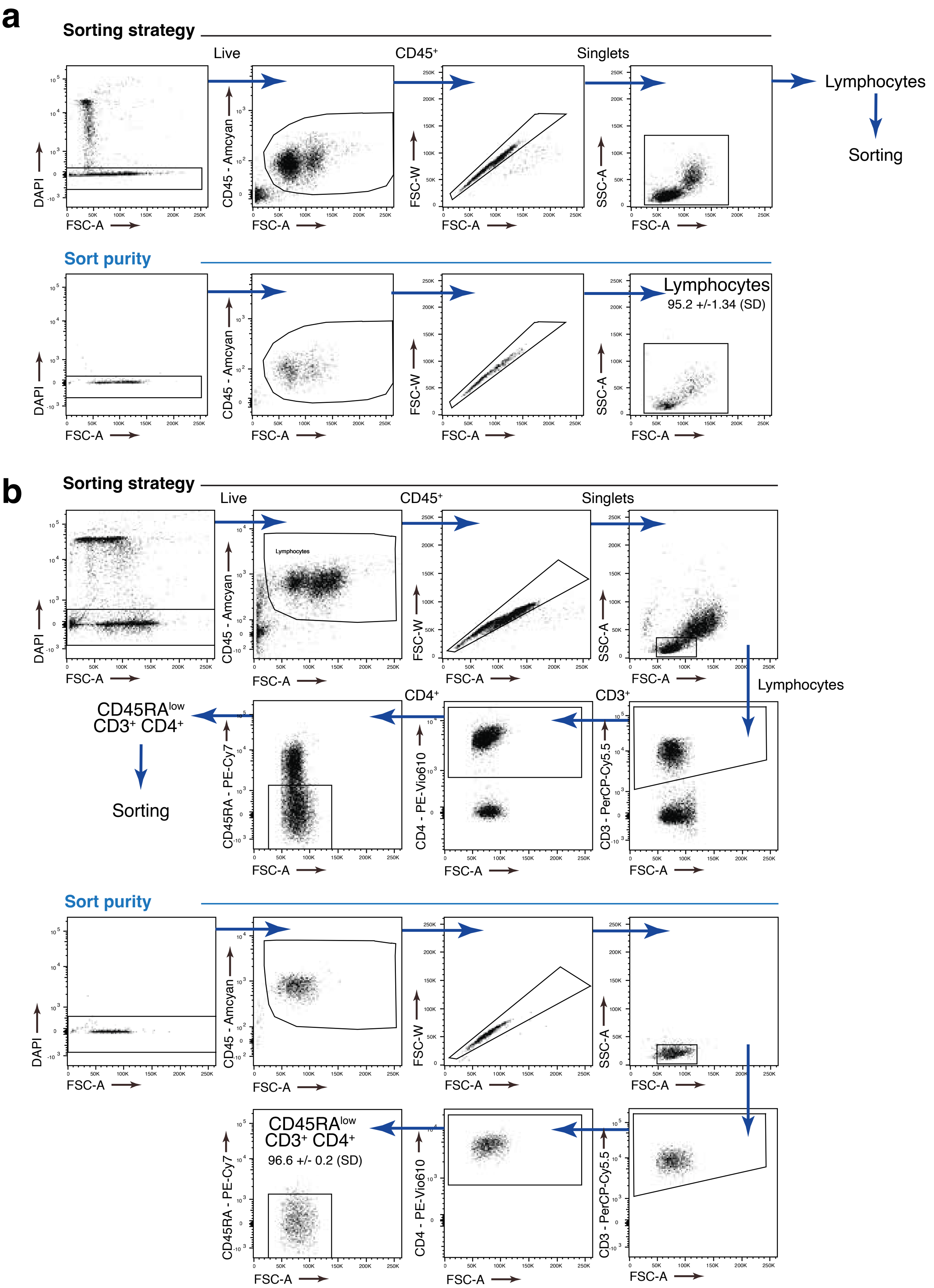


Technical Supplementary Figure 6

**Technical Supplementary Fig. 6. Fluorescence-activated cell sorting of peripheral blood mononuclear cells and $CD4^+$ memory T cells for single-cell sequencing.**

(a) Sorting strategy of PBMCs (upper panel) and purity control after sorting (lower panel) are shown.

(b) Sorting strategy of $CD4^+$ memory T cells (upper panel) and purity control after sorting are shown.

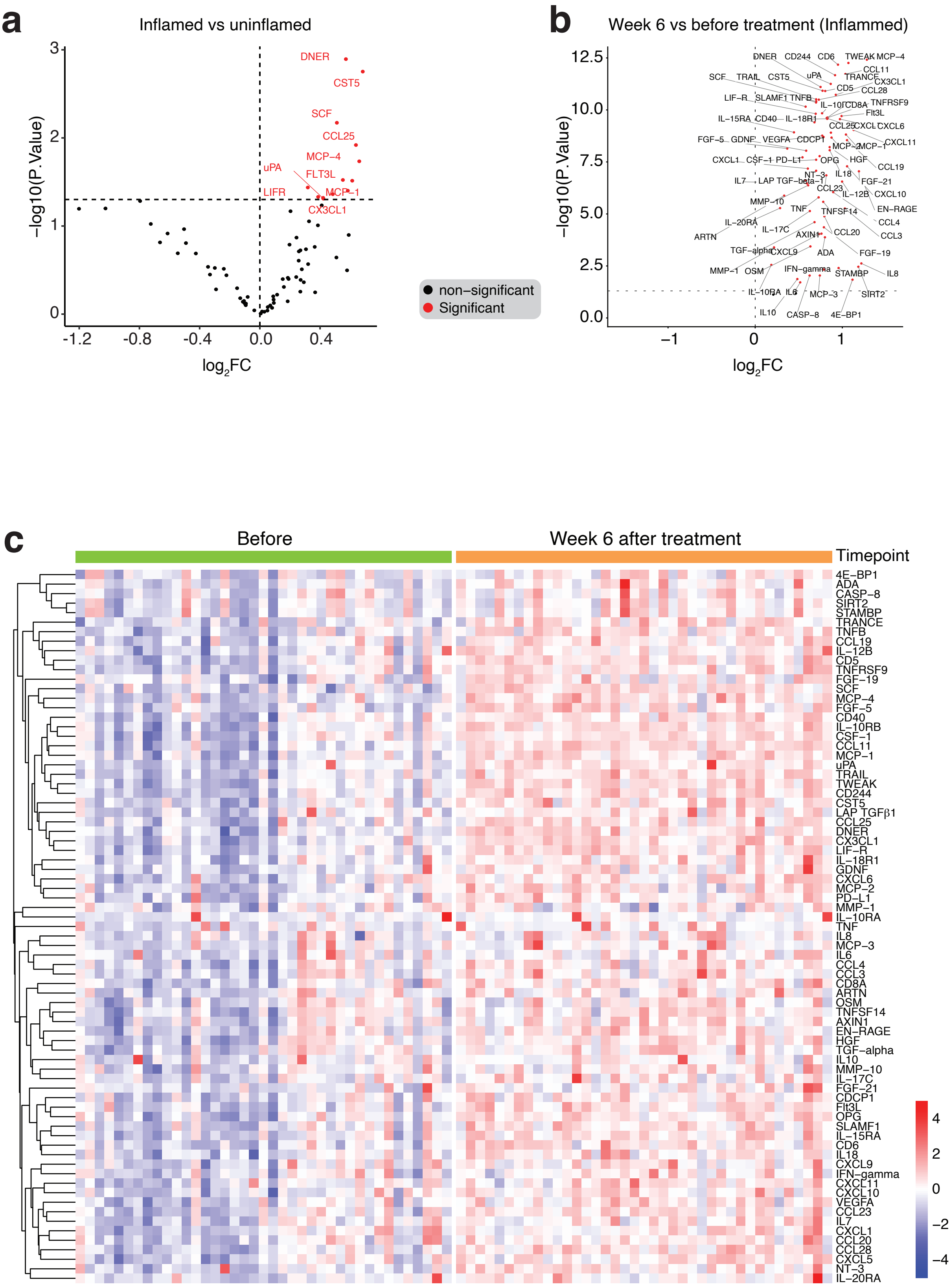


Technical Supplementary Figure 7

**Technical Supplementary Fig. 7. Serum proteomics from healthy control and IBD patients.**

(a) Volcano plot showing differential expression of serum protein concentrations (NPX values) between HD vs. IBD patients before therapy

(b) Volcano plot showing differential expression of serum protein concentrations in IBD patients before and 6 weeks after treatment with vedolizumab.

(c) Heatmap of serum proteins in IBD patients before (week 0) and after treatment.

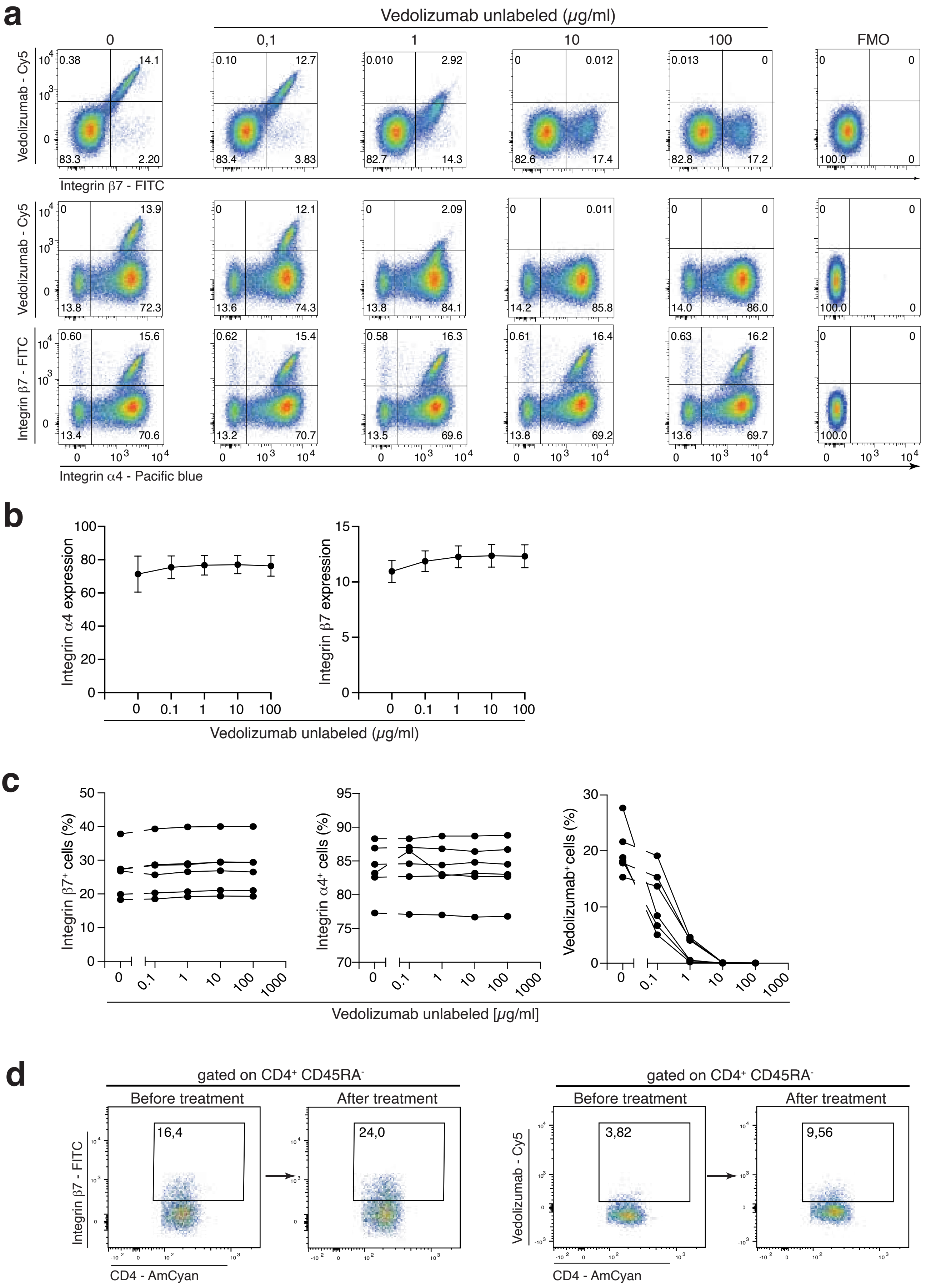


Technical Supplementary Figure 8

**Technical Supplementary Fig. 8. Impact of vedolizumab treatment on the detection of integrin α4β7 by flow and mass cytometry.**

(a-c) Staining of integrin α4 and integrin β7 after *in vitro* treatment of PBMCS with vedolizumab. PBMCs were isolated and incubated with unlabelled vedolizumab with the indicated concentrations. Integrin α4β7 was detected by Cy5-labelled vedolizumab (DRFZ) or FITC-labelled anti-integrin β7 antibody (clone FIB504). (b) The expression intensity (Geo mean index, expression normalized to naïve $CD4^+$ T cells) of integrin α4 and integrin β7 is shown. (c) Frequency of integrin $\alpha4^+$ and integrin $\beta7^+$ $CD4^+$ memory T cells are shown after treatment with vedolizumab with the indicated concentrations. Data representative of 3 experiments with a total n=6.

(d) Integrin $\beta7^+$ staining on SMART-tube fixed whole blood from an IBD patient before (0 weeks) and after (6 weeks) of treatment. A representative donor is shown.
