## Supplementary material for "Multimodal profiling of peripheral blood identifies proliferating circulating effector CD4^+^ T cells as predictors for response to integrin α4β7-blocking therapy in patients with inflammatory bowel disease": Suppl. Table 2

**Suppl. Table 2. Parameters used for machine learning.**

| Flow Cytometry | Olink | CyTOF | Clinical parameters |
|---|---|---|---|
| RorgT$^+$ Tbet$^+$ (memory CD4 T cells) | IL8 | Basophils | Leukocytes |
| EOMES (memory CD4 T cells) | VEGFA | cDCs | CRP |
| GATA3 (memory CD4 T cells) | CD8A | cMonocytes | Thrombocytes |
| RorgT (memory CD4 T cells) | MCP-3 | Eosinophils | Hemoglobin |
| T-bet (memory CD4 T cells) | GDNF | gd T cells | Neutrophils |
| Treg (CD4 T cells) | CDCP1 | Memory B cells | |
| IFN-γ (memory CD4 T cells) | CD244 | Naive B cells | |
| IL-10 (memory CD4 T cells) | IL7 | MAIT cells | |
| IL-17A. (memory CD4 T cells) | OPG | Memory CD4 | |
| IL-22 (memory CD4 T cells) | LAP TGF-beta-1 | Memory CD8 | |
| IL-4 (memory CD4 T cells) | uPA | Naive CD4 | |
| TNF-α (memory CD4 T cells) | IL6 | Naive CD8 | |
| IL-10_Treg (CD4 T cells) | IL-17C | ncMonocytes | |
| Integrin α4 (memory CD4 T cells) | MCP-1 | NK cells | |
| Integrin β1 (memory CD4 T cells) | IL-17A | pDCs | |
| Integrin β7 (memory CD4 T cells) | CXCL11 | Plasmablasts | |
| CCR6 (memory CD4 T cells) | AXIN1 | | |
| CCR7 (memory CD4 T cells) | TRAIL | | |
| CCR9 (memory CD4 T cells) | CXCL9 | | |
| CD25 (memory CD4 T cells) | CST5 | | |
| CD38 (memory CD4 T cells) | OSM | | |
| CD103 (memory CD4 T cells) | CXCL1 | | |
| CD127 (memory CD4 T cells) | CCL4 | | |
| CD161 (memory CD4 T cells) | CD6 | | |
| CTLA4 (memory CD4 T cells) | SCF | | |
| CXCR3 (memory CD4 T cells) | IL18 | | |
| GPR15 (memory CD4 T cells) | SLAMF1 | | |
| HLADR (memory CD4 T cells) | TGF-alpha | | |
| Ki67 (memory CD4 T cells) | MCP-4 | | |
| PD1 (memory CD4 T cells) | CCL11 | | |
| HLA-DR$^+$ CD38$^+$ (memory CD4 T cells) | TNFSF14 | | |
| | IL-10RA | | |
| | MMP-1 | | |
| | LIF-R | | |
| | FGF-21 | | |
| | CCL19 | | |
| | IL-15RA | | |
| | IL-10RB | | |
| | IL-18R1 | | |
| | PD-L1 | | |
| | CXCL5 | | |
| | TRANCE | | |
| | HGF | | |
| | IL-12B | | |

| | |
|---|---|
| | MMP-10 |
| | IL10 |
| | TNF |
| | CCL23 |
| | CD5 |
| | CCL3 |
| | Flt3L |
| | CXCL6 |
| | CXCL10 |
| | 4E-BP1 |
| | SIRT2 |
| | CCL28 |
| | DNER |
| | EN-RAGE |
| | CD40 |
| | IFN-gamma |
| | FGF-19 |
| | MCP-2 |
| | CASP-8 |
| | CCL25 |
| | CX3CL1 |
| | TNFRSF9 |
| | NT-3 |
| | TWEAK |
| | CCL20 |
| | ST1A1 |
| | STAMBP |
| | ADA |
| | TNFB |
| | CSF-1 |
