## Supplementary material for "Multimodal profiling of peripheral blood identifies proliferating circulating effector CD4^+^ T cells as predictors for response to integrin α4β7-blocking therapy in patients with inflammatory bowel disease": Suppl. Table 5

**Suppl. Table 5. Cohort 2 - Patient characteristics employed in this study (Figure 1i).**

| | | **Ulcerative colitis** |
|---|---|---|
| **Number of patients** | | 15 |
| **Age (mean, range)** | | 42 (22-75) |
| **Female (%)** | | 20% |
| **HBI (mean, range)** | | |
| **Partial Mayo Score (mean, range)** | | 4,2 (1-7) |
| **Co-medication (%)** | None | 46,67% |
| | Prednisolon | 26,67% |
| | Budenofalk | 13,33% |
| | Azathioprin | 6,67% |
| | Prednisolon + Budenofalk | 6,67% |
| **Disease localisation** | | E1 (proctitis):33,33% |
| | | E2 (left sided): 33,33% |
| | | E3 (extensive): 33,33% |
