## Supplementary material for "Multimodal profiling of peripheral blood identifies proliferating circulating effector CD4^+^ T cells as predictors for response to integrin α4β7-blocking therapy in patients with inflammatory bowel disease": Suppl. Table 7

**Suppl. Table 7. Patient characteristics used for flow cytometry.**

| | | Healthy control | Crohn's disease | Ulcerative colitis |
|---|---|---|---|---|
| **Number of patients** | | 26 | 9 | 32 |
| **Age (mean, range)** | | 41,04 (22-75) | 44,78 (23-75) | 40,66 (19-74) |
| **Female (%)** | | 42,31% | 88,9% | 59,38% |
| **HBI (mean, range)** | | | 9,44 (4-27) | |
| **Partial Mayo Score (mean, range)** | | | | 2,78 (0-7) |
| **Co-medication (%)** | None | | 55,56% | 59,38% |
| | Prednisolon | | 22,22% | 31,25% |
| | Azathioprin | | 22,22% | 3,13% |
| | Budenofalk | | | 3,13% |
| | Prednisolon+ Cyclosporin | | | 3,13% |
| **Disease localisation** | | | L1 (ileal): 22,22% | |
| | | | L2 (ileocolonic): 11,11% | E1 (proctitis): 12,5% |
| | | | L3 (colonic): 22,22% | E2 (left sided): 46,88% |
| | | | L1/L2 (ileocolonic): 22,22% | E3 (extensive): 40,63% |
| | | | L1/L4: 11,11% | |
| | | | L3/4 (colonic): 11,11% | |
