## Supplementary material for "Multimodal profiling of peripheral blood identifies proliferating circulating effector CD4^+^ T cells as predictors for response to integrin α4β7-blocking therapy in patients with inflammatory bowel disease": Suppl. Table 9

**Suppl. Table 9. Patient characteristics used for mass cytometry.**

| | | Healthy control | Crohn's disease | Ulcerative colitis |
|---|---|---|---|---|
| **Number of patients** | | 26 | 9 | 22 |
| **Age (mean, range)** | | 40,23 (22-75) | 44,78 (23-75) | 43,82 (19-77) |
| **Female (%)** | | 61,5 % | 88,9% | 63,63% |
| **HBI (Ø, range)** | | | 9,44 (4-27) | |
| **Partial Mayo Score (Ø, range)** | | | | 3 (0-6) |
| **Co-medication (%)** | None | | 55,56% | 40,9% |
| | Prednisolon | | 22,22% | 31,81% |
| | Azathioprin | | 22,22% | 4,55% |
| | Infliximab | | | 4,55% |
| | Prednisolon+ Cyclosporin | | | 9,1% |
| | Budenofalk | | | 4,55% |
| **Disease localisation (%)** | | | L1 (ileal): 22,22% | E1 (proctitis):22,72% |
| | | | L2 (ileocolonic): 11,11% | E2 (left sided): 36,36% |
| | | | L3 (colonic): 22,22% | E3 (extensive): 40,91% |
| | | | L1/L2 (ileocolonic): 22,22% | |
| | | | L1/L4: 11,11% | |
| | | | L3/4 (colonic): 11,11% | |
