## Supplementary material for "Multimodal profiling of peripheral blood identifies proliferating circulating effector CD4^+^ T cells as predictors for response to integrin α4β7-blocking therapy in patients with inflammatory bowel disease": Suppl. Table 11

**Suppl. Table 11. Patient characteristics used for CITE-seq analysis.**

| | | Healthy control | Ulcerative colitis |
|---|---|---|---|
| **Number of patients** | | 5 | 10 |
| **Age (mean, range)** | | 34,6 (22-60) | 46,1 (19-77) |
| **Female (%)** | | 60% | 70% |
| **Partial Mayo Score (mean, range)** | | | 3,1 (0-6) |
| **Co-medication (%)** | None | | 30% |
| | Prednisolon | | 40% |
| | Infliximab | | 10% |
| | Prednisolon+ Cyclosporin | | 20% |
| **Disease localisation (%)** | | | E1 (proctitis):20% |
| | | | E2 (left sided): 40% |
| | | | E3 (extensive): 40% |
