## Supplementary material for "Multimodal profiling of peripheral blood identifies proliferating circulating effector CD4^+^ T cells as predictors for response to integrin α4β7-blocking therapy in patients with inflammatory bowel disease": Suppl. Table 12

**Suppl. Table 12. Patient characteristics used for Olink analysis.**

| | | Healthy control | Crohn's disease | Ulcerative colitis |
|---|---|---|---|---|
| **Number Patients** | | 12 | 7 | 31 |
| **Age (mean, range)** | | 46,58 (22-75) | 48,29 (23-75) | 40,97 (19-77) |
| **Female (%)** | | 58,33% | 85,71% | 46,88% |
| **HBI (mean, range)** | | | 10,14 (4-27) | |
| **Partial Mayo Score (mean, range)** | | | | 2,78 (0-7) |
| **Co-medication (%)** | None | | 42,85% | 56,25% |
| | Prednisolon | | 28,57% | 31,25% |
| | Azathioprin | | 28,57% | 3,13% |
| | Infliximab | | | 3,13% |
| | Prednisolon+ Cyclosporin | | | 6,25% |
| **Disease localisation** | | | L2 (ileocolonic): 14,28% | E1 (proctitis):15,63 % |
| | | | L3 (colonic): 28,57% | E2 (left sided): 43,75% |
| | | | L1/L2 (ileocolonic): 28,57% | E3 (extensive): 40,63% |
| | | | L1/L4: 14,28% | |
| | | | L3/4 (colonic): 14,28% | |
